# Flattening the curve - How to get better results with small deep-mutational-scanning datasets

**DOI:** 10.1101/2023.03.27.534314

**Authors:** Gregor Wirnsberger, Iva Pritišanac, Gustav Oberdorfer, Karl Gruber

## Abstract

Proteins are utilized in various biotechnological applications, often requiring the optimization of protein properties by introducing specific amino acid exchanges. Deep mutational scanning (DMS) is an effective high-throughput method for evaluating the effects of these exchanges on protein function. DMS data can then inform the training of a neural network to predict the impact of mutations. Most approaches employ some representation of the protein sequence for training and prediction. As proteins are characterized by complex structures and intricate residue interaction networks, directly providing structural information as input reduces the need to learn these features from the data.

We introduce a method for encoding protein structures as stacked 2D contact maps, which capture residue interactions, their evolutionary conservation, and mutation-induced interaction changes. Furthermore, we explored techniques to augment neural network training performance on smaller DMS datasets. To validate our approach, we trained three neural network architectures originally used for image analysis on three DMS datasets, and we compared their performances with networks trained solely on protein sequences. The results confirm the effectiveness of the protein structure encoding in machine learning efforts on DMS data. Using structural representations as direct input to the networks, along with data augmentation and pre-training, significantly reduced demands on training data size and improved prediction performance, especially on smaller datasets, while performance on large datasets was on par with state-of-the-art sequence convolutional neural networks.

The methods presented here have the potential to provide the same workflow as DMS without the experimental and financial burden of testing thousands of mutants. Additionally, we present an open-source, user-friendly software tool to make these data analysis techniques accessible, particularly to biotechnology and protein engineering researchers who wish to apply them to their mutagenesis data.

## 3 Introduction

Proteins are found in viruses, bacteria, plants, and humans and fulfill a huge number of different functions and tasks in living organisms. Given their enormous functional diversity, proteins also present an attractive platform for various applications in biotechnology and bioengineering. However, naturally occurring proteins often require optimization for non-native uses. One common method of protein optimization involves the substitution of specific amino acids, which can significantly enhance or alter the protein’s function as, for instance, observed in the increased brightness of fluorescent proteins [1], or in antibody binding target modifications [2].

Amino acid substitutions can profoundly affect the properties of proteins, with mutagenesis providing a potent tool for evaluating these effects. A powerful technique for gaining comprehensive insights into genotype-phenotype relationships is deep mutational scanning (DMS) [3]. This approach enables the creation of expansive datasets depicting the effects of mutations on a given protein. DMS combines some type of protein display, which provides a physical link between a protein and its encoding nucleic acid sequence, with high-throughput sequencing, allowing for the characterization of up to 10^5^ protein variants. The methodology involves applying selective pressure based on the protein’s function to a diverse library of protein variants, which are sequenced before and after selection. High-throughput sequencing then quantifies the abundance of each variant. Throughout selection, variants with beneficial mutations become enriched, while those with deleterious mutations become depleted, offering a means to quantify the fitness of a vast sequence diversity for a protein of interest [4]. The broad applicability of DMS is demonstrated in its diverse uses, such as investigating the sequence determinants of A*β* aggregation in Alzheimer’s disease [5], probing protein binding behavior [6], forecasting the evolutionary trajectories of human H3N2 influenza variants [7], optimizing antimicrobial peptides [8], and elucidating the effects of mutations in SARS-CoV-2 proteins [9] [10].

DMS experiments have increasingly become the method of choice for many projects aiming to achieve specific engineering goals. As these experiments grow, there is an increasing demand for user-friendly predictive methods tailored to this kind of data. Consequently, various methods have been developed to predict the effects of amino acid exchanges in proteins. Some of these methods rely solely on evolutionary data and omit experimentally determined data to predict the functional consequences of amino acid substitutions. These approaches include, for example, the use of Hidden-Markov models [11], Potts models (EVmutation [12]), and variational autocoders (DeepSequence [13]). Others are natural language processing models, which are strongly influenced by the training approaches used in their field of origin. They get pre-trained in an unsupervised manner on a large amount of data and then fine-tuned on the prediction task. Here, models like LSTMs [14] and transformer [15] are used.

Additionally, some models employ decision tree ensembles (like Envision [16]) trained on deep mutational scanning data or use Gaussian processes [17] for predictions. These models, particularly those grounded in natural language processing (NLP), often take only the protein sequence as input. Other models, such as Envision, integrate structural features into their framework but tend to utilize more general features like secondary structures and solvent accessibility instead of harnessing the unique information that each amino acid can offer.

Another important aspect in training ML models is training efficient encoding of the underlying data. In the case of proteins, this can be the amino acid sequence alone without any 3D information [18], a graph representation of the protein structure [18], or voxel-based spatial structural encoding [19]. In recent years, models used in natural language processing have increasingly been applied to problems with proteins. Although these models are compelling and can produce great results, they tend to need a massive number of parameters, leading to high memory and computation requirements [20].

Since protein structure is more conserved than sequence [21], we created a - to our knowledge - new encoding for protein structures to take advantage of the information contained in the 3D structure. The encoding consists of 2D contact maps representing different physico-chemical properties of amino acids and their accompanying interaction, as well as the evolutionary conservation of each interacting residue in the structure (Section 4.2). In addition, this encoding allows the use of standard architectures for image classification networks, thus giving access to a large number of different architectures that can be used to solve this problem. Furthermore, we create a helpful pre-training and data augmentation protocol that helps to improve results when only a small amount of data is available (Fig. 1).

**Figure 1:**
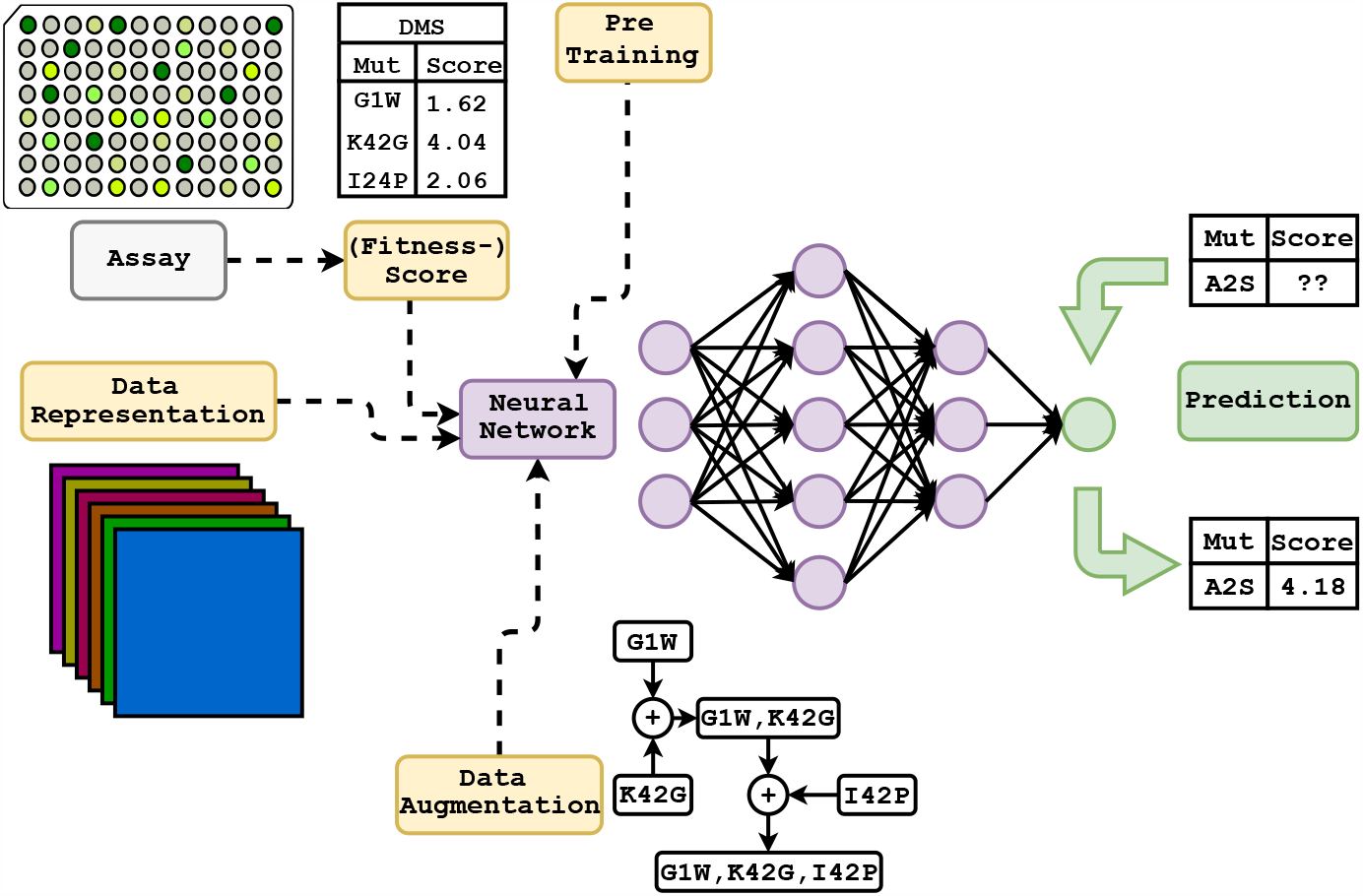
Overview of the training and prediction workflow. Initially, models are pre-trained on predicting a pseudo score that arises from the data representation (consisting of stacked 2D contact maps representing different physico-chemical properties and evolutionary information). This helps the model adjust its weights to the kind of prediction it will later be used for while not requiring additional data acquisition. Data augmentation is applied to up-size the training data to improve the prediction quality further. This is then used to train the network on experimentally determined (fitness-) scores of the protein of interest. In the end, the trained model can be used to predict these scores and, therefore, the effects of amino acid exchanges in the protein that were not experimentally determined. It is also possible to omit pre-training and data augmentation and train the network solely on experimentally determined data. Three different network architectures were used in this study, but they can be easily changed to any architecture of choice that accepts the input in the form of the data representation.

In order to determine the effectiveness of our approach when training data is scarce, we trained different architectures using datasets containing between 50 and 6000 samples. To ensure the accuracy of our analysis, we used sub-datasets that accurately reflect the distribution of fitness scores present in the complete datasets. This allowed us to determine the number of lab-tested variants required as training data to reliably model the underlying fitness score distribution of a protein’s fitness landscape. Additionally, we tested the ability of the networks to predict the effect of amino acid exchange on protein sequence positions that were not included in the training data. To further evaluate how the networks can cope with limited data from traditional mutation experiments, we trained models using data from a simulated extended alanine scan.

We assessed the performance of the same architecture with both sequence input and our structure representation, as well as the impact of pre-training and data augmentation. For this purpose, we relied on a recently published study by Gelman *et al*. [18]. Their work offers a comprehensive analysis of DMS datasets and evaluates the applicability of networks trained with sequence input on large datasets. Since these authors employed a simpler convolutional neural network architecture (CNN), we were able to use the same network architecture for our approach, enabling comparisons that are not influenced by architecture complexity or the use of distinct neural network architectures. This also facilitated comparisons with more complex CNN architectures and their potential benefits.

We further examined the performance of architectures with fewer parameters, revealing that while our representation provides an advantage, data augmentation and pre-training are crucial for optimal performance. Our workflow also demonstrated robust performance with an architecture that significantly reduces the number of parameters.

To promote the utilization of these methods in biotechnology and protein engineering, we provide open-source software featuring a user-friendly command line interface designed to be accessible to non-ML experts. Executing the program with new DMS data requires minimal input, but the software also provides numerous advanced settings if needed in specific cases.

## 4 Materials and methods

### 4.1 Data

In our study, we utilized DMS data previously prepared and used in the study by Gelman *et al*. [18]. We specifically chose data from avGFP, Pab1, and GB1, as these proteins demonstrated the best results in their study, making them ideal for comparison as the data set quality does not influence the results. As Gelman *et al*. [18] already explored the influence of data quality on learning performance, finding a strong correlation between predictive performance and data quality, we opted to use these three high-quality datasets and then tested the influence of dataset size, pre-training, data augmentation, encoding, and network architecture. A limited analysis, which included only the optimal settings and the biggest and smallest of the three architectures used, was performed on two lower-quality datasets (Bgl3 and Ube4b) also used in [18], where they exhibited poorer performance. Regarding protein structures, we also relied on data used in [18] to ensure the sequence, and consequently, the structure matched the DMS data. Therefore, we used the PDB files of these structures provided in the corresponding GitHub repository [22].

The DMS datasets also contain nonsense mutations. We chose not to use assay scores for proteins featuring one or more nonsense mutations since these scores would represent protein fragments and thus would not reflect the properties of the wild-type protein containing a particular mutation. We, therefore, modified the datasets to exclude all nonsense mutations during training, validation, and testing.

### 4.2 Interactions and their encoding

To emulate the effect of different mutations in a protein, we created interaction matrices that used a set of different amino acid properties to describe the interactions between residues in a protein and their changes due to amino acid exchanges. Additionally, a matrix that encoded the evolutionary conservation of interacting residues and an index matrix were used. Visual representations of the individual matrices, using Pab1 as an example, can be found in Fig. S4. This encoding method relies on the availability of the complete structure of the protein. In real-world scenarios, experimental structural data might not always be available or complete. However, there are a variety of approaches to address this issue, such as filling missing loops or even using advanced protein structure prediction tools like AlphaFold [23] to model the entire protein structure. In a worst-case scenario, in which only incomplete structures are available, the encoding can still work but would require dataset modifications (*e*.*g*., index adjustments based on the missing residues).

#### 4.2.1 Distance Matrix

To classify pairs of residues as interacting, we used Euclidean distances (*d*_*ij*_) calculated from Cartesian coordinates of all protein atoms stored in the corresponding PDB file [24]. Interacting residues were identified by checking the closest distance between side chain atoms of two residues, *i* and *j*. Using this approach, the smallest distances between all residues were calculated, and a symmetric *n × n* distance matrix (D), where *n* denotes the sequence length, was generated. Using equation 1, this matrix was then used to generate a so-called factor matrix (F).

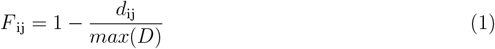

In Eq: 1 *d* _ij_ denotes the distance between two residues and *max(D)* the biggest distance seen in the structure.

This factor matrix was used to scale the “strength” of the interactions in all subsequent matrices (apart from the position matrix (P)) by calculating the Hadamard product (element-wise product) of F with each interaction matrix. Elements in F corresponding to distances larger than 20 Å were set to zero. This led to higher values for close interactions and smaller ones for interactions of residues that are further apart. In addition, it masked interactions originating from residues further apart than 20 Å.

#### 4.2.2 Index Matrix

Convolution neural networks (CNN) are translation invariant. This is one of the features that make them powerful in image recognition tasks since they can find patterns they have learned anywhere in an image and not rely on their position. In our case, this translation invariance was an undesirable feature because the positions of the interactions matter. To address this issue, we introduced a simple position matrix (P). It describes the position of each interaction in the matrices based on the index matrix I (Eq: 2). To calculate P, the Hadamard product of D and I is formed where D is set to 1 for distances smaller than dist_th_ and to 0 for bigger distances.

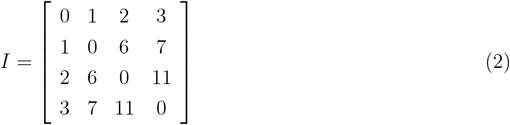

#### 4.2.3 Hydrogen Bonding

The number of hydrogen bonds is one of the factors that determine the stability of a protein. Therefore it is a crucial kind of interaction since amino acid exchanges that introduce hydrogen bonding capabilities or remove them will thus alter this property. Not all amino acids have the same capability of forming hydrogen bonds with their side chain. Some can only act as a donor (K, R, W), some as an acceptor (D, E), some as donors or acceptors (H, N, Q, S, T, Y), and some are not able to form hydrogen bonds with their side chain at all (A, C, F, G, I, L, M, P, V). The hydrogen bonding matrix B features a value of 1 for interactions formed by a donor and an acceptor, by a donor and an acceptor/donor, by an acceptor acceptor/donor or by an acceptor/donor acceptor/donor pair, or a value of 0 otherwise.

#### 4.3.4 Hydrophobicity

Proteins often contain a hydrophobic core and a hydrophilic outside that interacts with its surroundings. The hydrophobic core plays an important role in the folding process of a protein. Therefore, mutations that change the hydrophobicity in certain areas of a protein can have positive and negative effects. The hydrophobicity values used were obtained from Parrot [25]. These hydrophobicity values range from -4.5 for arginine to 4.5 for isoleucine.

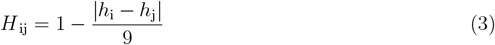

In Eq: 3 *h* denotes the hydrophobicity of a certain residue, and 9 is the maximum possible hydrophobicity difference. The hydrophobicity matrix (H) describes how well-interacting residues match in terms of their hydrophobicity.

#### 4.2.5 Charge

There are three main types of amino acids categorized according to their charge: neutral (A, C, F, G, I, L, M, N, P, Q, S, T, W, Y), positively charged (R, H, K), and negatively charged (D, E). Salt bridges, which are interactions of residues of opposite charge, are, besides hydrogen bonds, another type of interaction that is important for the stability of a protein. On the other hand, amino acids that carry the same charge can repel each other, which can lead to instability in the protein’s structure. To calculate the charge matrix (C) (where we multiply the amino acids charge value and this result by -1), we assigned a value of 1 to interactions between positively charged amino acids, a value of -1 to interactions between amino acids carrying the same charge, and a value of 0 to all other interactions.

#### 4.2.6 Surface accessible side chain area

Amino acids feature a variety of different sizes of their side chain. This is reflected in the difference in their surface accessible side chain area (SASA). The bigger the SASA, the higher the possibility for a (strong) interaction. Therefore a mutation that changes the interaction area between two interacting residues can have an influence on their interaction strength. The SASA values were obtained from Parrot [25], ranging from 0 Å^2^ for glycine to 254 Å^2^ for tryptophan.

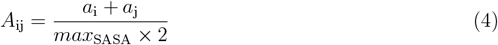

In Eq: 4.2 *a* denotes the interaction area of a certain residue and *max* _SASA_ the maximum SASA value for an amino acid. The interaction area matrix (A) describes the interaction area between residues.

#### 4.2.7 Clashes

Amino acids also differ in the length of their side chains. That means certain mutations can lead to potential “holes” in a protein if the side chains get shorter or potential clashes because the side chains are too long for the space between them.

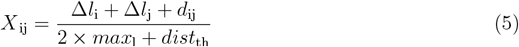

In Eq: 5 **Δ***l* denotes the change in the side chain length at a certain residue position from wild-type to the variant, *max* _l_ the maximum side chain length and *dist*_th_ the maximum allowed distance between two residues to count as interacting. Side chain lengths range from 0 Å for glycine to 8.28 Å for arginine. To obtain the values of the side chain length, we used Pymol [26] to measure the maximum distances between the C*α* and side chain atoms in different residue types. The resulting clash matrix (X), represented by Eq: 5, shows the distances between interacting side chain residues. If a mutation leads to a distance between two residues that is closer than the distance between them in the wild-type, a negative length value is recorded. This means that the values in this matrix, along with the charge matrix C, are the only ones that fall within the range of [-1, 1] instead of [0, 1].

#### 4.2.8 Evolution

To make use of the evolutionary information that can be obtained through a Blast search [27], we create a matrix (E) based on the conservation of amino acids at each sequence position. Therefore we used the result of a blastp search against the wild-type protein sequence with its default settings against the experimental clustered non-redundant database and aligned the obtained sequences as well as the wild-type sequence using the multiple sequence alignment tool Clustal omega [28]. Duplicated sequences were removed from the alignment. To calculate a conservation score at each wild-type sequence position, all present amino acids were counted at this position, and their counts were divided by the total number of amino acids present at that position. Amino acids that were not present at this position got a value of zero assigned. To evaluate the conservation of an interaction, the conservation scores of the interacting residues were multiplied. Evolutionary information could also be integrated via, e.g., a separate branch of the neural network, but we chose this representation because it was easier to incorporate into the existing network structure. Additionally, this representation encodes the change in the conservation of an interaction based on the exchanged amino acid(s).

Figure S4 shows an example of all interaction matrices (B, H, C, A, X) for Pab1 containing the mutation “N127R, A178H, G177S, A178G, G188H, E195K, L133M, P125S” as well as the position matrix (P), the interaction matrix (M) which describes which residues interact with each other and the distance matrix (D).

### 4.3 Network architectures

#### 4.3.1 Simple CNN

Since we wanted to compare our structure representation to the sequence convolution approach (Section: 4.3.4), a LeNet5 [29] - like convolutional neural network (Fig. S5) was used. It contains a feature extraction part containing three 2D convolution layers with 16, 32, and 64 filters and a kernel size of 3×3, each followed by a max pooling layer. After that comes a flatten layer and a classifier part consisting of 4 fully connected layers with 128, 256, 256, and 64 nodes and a single output node. We used the leaky rectified linear unit (leaky RELU) as the activation function for all layers in the model. Zero-padding was used throughout the whole network. This model is referred to as “simple CNN”.

#### 4.3.2 DenseNet

To compare the performance to a more recently described architecture, we chose to use a DenseNet [30] - like architecture (Fig. S6), which will be referred to as “DenseNet”. Here the core building block consists of a 2D convolution layer with 128 filters and a kernel size of 1×1, followed by a 2D convolution layer with 12 filters and a kernel size of 3×3. Zero-padding is used throughout the whole network to keep each layer’s input and output dimensions the same. The input into the first 2D convolution layer and the output of the second get concatenated. This is repeated 4 (*block depth*) times and is then followed by a 2D average pooling layer with a kernel size of 2×2. All this combined is one block, and this is repeated 4 (*block number*) times. In the end, a 2D global average pooling layer is followed by a fully connected network with 128, 128, and 64 nodes per layer leading into one output node. Additionally, we used an “intro layer” for avGFP, which consists of a 2D convolution layer with 128 filters, a kernel size of 3×3, and a stride of 2 followed by a 2D max pooling layer with a kernel size of 3 and a stride of 2 at the beginning of the network. This reduces the size of the input and thereby reduces the number of computations needed in the rest of the network. In contrast to the original DenseNet, we omitted batch normalization because it led to worse performance and used the leaky RELU instead of RELU as the activation function.

#### 4.3.3 SepConvMixer

To test the performance of a network with as few parameters as possible, we implemented an architecture (Fig. S7) similar to ConvMixer [31]. Sequence convolution requires up to 82 times, simple CNN up 185 times, and DenseNet up to 21 times the number of parameters in our settings (Table 4). The two main contributors to the reduction of the number of parameters were the possibility of using a smaller fully connected classifier network as well as the use of 2D separable convolution layers. The latter first performs a depth-wise spatial convolution, which acts separately on each input channel and is followed by a point-wise convolution to mix the resulting output channels. The network starts with one 2D separable convolution layer with 32 filters where we used a kernel size of 3×3 and a stride of 1 for smaller proteins (like Pab1 and GB1) and a kernel size of 9×9 and a stride of 9 for bigger proteins (avGFP). This is followed by a variable number of blocks (determined by the parameter *depth*) each consisting of 2 2D separable convolution layers with 32 filters and a kernel size of 3×3. The input into the first, the output of the first, and the output of the second layer get added at the end of the block. We used a *depth* of 9 in this study. These blocks are followed by a 2D global max pooling layer and a fully connected network consisting of 128- and 64-node layers followed by a single-node output layer. We used the leaky RELU as well as zero-padding to keep the dimensions the same throughout the whole network.

A “down-sampling” (a kernel size of 9×9 with a stride of 9 in the first layer) for bigger proteins slightly reduces the performance but is a worthy trade-off to reduce the computational cost.

#### 4.3.4 Sequence convolution

For comparison, we used the network architectures of [18] as specified in their main experiments (/pub/regression args/PROTEIN main cnn.txt [22]). Apart from enabling early stopping and restricting the length of the training to 100 epochs, we chose the default parameters when using the /code/regression.py. This is referred to as “sequence convolution” throughout the paper.

#### 4.3.5 Implementation

Our models were implemented using Python v3.10, TensorFlow v2.9.1, and Keras v2.9.0

### 4.4 Training

Training of simple CNN, DenseNet, and SepConvMixer architectures was performed using the mean absolute error as the metric, Adam as optimizer with a learning rate of 0.001 and a maximum number of epochs of 100. Furthermore, we stopped the training if the mean absolute error did not improve by at least 0.01 over 20 epochs. The batch size for the training was 32 and parallelized by using 12 central processing unit (CPU) cores of an AMD Ryzen Threadripper 3960X. The training was performed on an Nvidia RTX A5000 graphics processing unit (GPU). For training the networks on the ANH-Scan data, an Nvidia GeForce RTX 3070 and an Intel Xeon Gold 6230R CPU were used. For pre-training, we limited the maximum number of epochs to 70. The training of the sequence convolution network was done using an Intel Xeon Gold 6230R CPU.

### 4.5 Experiment setup

#### 4.5.1 Dataset size effect

Data and dataset selection can have an impact on the performance of the neural network. To avoid any advantage or bias through the use of only specific subsections of the data, *e*.*g*., only low DMS scores, we selected the training, validation, and test dataset in the following way: The whole dataset was randomly shuffled. The first *n* data points were used as training data, the following *n ×* 0.2 samples were used as validation data during the training, and the next 5000 data points were used as test dataset after the training, where *n* is the training data size. This ensures that the training-, validation- and test datasets are entirely disjoint and do not feature overlapping data. Since the artificially created pre-training data has a Pearson correlation of around -0.5 to the DMS data, the pre-training datasets were created so that the data points in the pre-training dataset do not feature mutations that are in the test dataset to ensure no knowledge leak and an unbiased test result.

This led to training- and test datasets that featured a similar DMS score distribution as the whole dataset and, therefore, built a representative sample (Figure: S1 - S3). For each training run, we used three different data sets, which were all obtained from the original data sets of the proteins: a train, a tune, and a test set. The test set always consisted of 5000 randomly chosen unique entries each. The tune set had one-fifth of the size of the training data set for our architectures and always 5000 entries for sequence convolution. The train datasets contained 50, 100, 250, 500, 1000, 2000, or 6000 entries for all training runs. The train data set was used to train the network, the tune set was used to calculate the validation statistics during training, and the test set was used to calculate the statistics of the performance of the network after training. Training simple CNN, SepConvMixer, DenseNet, and sequence convolution was done on three randomly chosen subsets of the whole protein data sets to construct the train, tune, and test sets to avoid picking one that suits one architecture better by chance.

For the training of simple CNN, SepConvMixer, and DenseNet, we used data augmentation (Section: 4.5.1) as described below, as well as pre-training (Section: 4.5.1). For sequence convolution, we used the same train, tune, and test sets as for the training of simple CNN, SepConvMixer, and DenseNet; however, we did not use data augmentation and transfer learning during its training process. Three main performance metrics are used: mean squared error (MSE), Pearson’s correlation coefficient, and Spearman’s correlation coefficient, with the main focus on Pearson’s correlation coefficient. No dedicated hyper-parameter tuning was done, but those that had proven to be the best after some initial testing were used. To test the impact of an “intro layer” like in the original DenseNet, which is a normal 2D convolution layer with a kernel size of 3×3 and a stride of 2 followed by a 2D max pooling layer with a kernel size of 3×3 and a stride of 2, we chose to include this in the training of avGFP but not for Pab1 and GB1. The same was done for SepConvMixer, where the first separable convolution layer has either a kernel size of 3 and a stride of 1 or, for avGFP, a kernel size of 9 and a stride of 9. The use of an “intro layer” reduced the performance for smaller proteins like Pab1 and GB1 slightly but is needed and a good trade-off to be computationally efficient for proteins of the size of avGFP and bigger.

## Data augmentation

Since neural networks learn better with more data, we used a simple data augmentation method to obtain more training data from small data sets. This method uses the given experimental data, e.g., Table 1, shuffles it, and adds it to the original not shuffled data to create new augmented variants like shown in Table 2.

**Table 1:** Sample data for augmentation.

| variant | number mutations | score |
| --- | --- | --- |
| K1L,S3A | 2 | -0.3 |
| R23H,W19F | 2 | 0.1 |
| C5G,A7L | 2 | -1.0 |

Shows data samples later used for an example of data augmentation.

**Table 2:** Sample augmentation.

| augmented variant | number mutations | augmented score |
| --- | --- | --- |
| K1L,S3A,R23H,W19F | 4 | -0.2 |
| R23H,W19F,C5G,A7L | 4 | -0.9 |
| C5G,A7L,K1L,S3A | 4 | -1.3 |

Shows how data points of Table 1 are added during data augmentation.

This is done four times, and the newly created data is stored. This data is then used as input data to perform the same action three times where, after each round, the newly created data is used as the new input data in the next round. From this newly created augmented data, as many samples are drawn as needed to get a maximum of 20000 training samples when the original data is added (*aug*_used_ = 20000 ***−*** *n*_original_ where aug_used_ is the number of augmented samples used and n_original_ the number of original data). If the augmentation does not produce enough data to reach a combined number of 20000 samples after the original data is added, the whole augmented data is used. It did not show good results when increasing the number of runs to produce +20000 samples when the original data set is not big enough to reach the number of samples with the number of runs described above. This kind of data augmentation produces pseudo labels for data and assumes an additive effect of mutations. Even though there are more intricate models to describe the relationship between different mutations in a protein, this method provides a simple and effective way to quickly generate more data that helps the model produce better results. In addition, the assumption of simple additivity does not rely on another model, such as DeepSequence [13], to be added to the training procedure. We also tried training the networks only on augmented data and fine-tuning them on the original small data sets. This showed worse performance than training them with the original and augmented data combined.

### Pre-training

To overcome the need for big data sets, we used pre-training to obtain better results while training on small data sets. The transfer of weights of the feature extraction part of a network trained on a whole dataset of another protein yields better performance than starting from a completely untrained network. However, to pre-train a network on data that is more closely correlated to the protein of interest, we created a pseudo-score that can be calculated without the need for experimental data (Section: 4.5.1). Since the pre-training is based on our encoding, we used it for simple CNN, SepConvMixer, and DenseNet. After training the model on the pseudo data, the weights of the feature extractor were transferred to an untrained network, frozen, and a new classifier was trained. The same was done with a trainable feature extractor. During initial tests, the reduction of the learning rate did not improve the performance. Therefore we omitted it in further studies. Transferring the weights of the whole pre-trained model, including the classifier, showed worse performance. We also tested networks pre-trained on other proteins, e.g., pre-trained on avGFP and trained on Pab1, but our pre-training method proved to be more effective.

### Pseudo Score

In order to calculate the pseudo score for the pre-training, the wild-type of the protein gets encoded in the same way as for training the network. The same is done for all possible single and double mutants of the protein. To calculate the pseudo score of a variant, the encoded wild type gets element-wise subtracted from the encoded variant matrix. Of all these values, the absolute value is taken and summed up over all matrices. This gets divided by 100 to shift the values into the range of the real fitness scores. 40000 of these created data points are randomly chosen and used to pre-train the models. These pseudo scores show a Pearson’s R of around -0.5 to the original DMS data for the different datasets.

#### 4.5.2 Positional extrapolation

To evaluate the networks’ capabilities to predict mutational effects of positions not seen during training, the protein sequence was divided into training and validation sets, comprising 85% of the positions, and a test set of the remaining 15%. This was done three times with randomly selected sequence positions. Multi-mutation variants with some positions in the test set and others in the training set were eliminated from this analysis. To test this, we used the pretrained networks (simple CNN and SepConvMixer) on our pseudo score from 4.5 and trained them on the data described above. To compare their performance, we also trained sequence convolution on the same data. We did this analysis for GB1, Pab1, and avGFP. The training dataset size for GB1 was 351000 data points, 23000 for Pab1, and 26000 for avGFP.

#### 4.5.3 ANH scan

An often method for assessing mutational effects in proteins is an alanine scan, where each amino acid is replaced with alanine and the property of interest is evaluated. This approach generates a limited dataset of the size equivalent to the length of the protein sequence. Recently, it has been discovered that the amino acid exchanges to alanine, asparagine, and histidine are the most correlated with all other single amino acid exchanges [32]. Therefore, to increase the amount of data and provide the neural network with a good starting point, an extension of the alanine scan was proposed, an ANH-scan [33]. In this regard, we selected from the DMS datasets all single variants that contain either an exchange to alanine, asparagine, or histidine as a training and validation dataset. 85% of these were used as training data, and 15% were used as validation data during training. The remaining single mutants of the datasets were used as test data. To test this approach, we used the networks (simple CNN and SepConvMixer) pre-trained on our pseudo score from 4.5.1 and trained them on the data described above. To compare their performance, we also trained the sequence convolution model on the same data. This approach yielded a combined train and tune dataset size of 159 for GB1, 132 for Pab1, and 169 for avGFP, which indicates that only for GB1 almost all positions were mutated to either A, N, or H and that the other data sets are missing some of these.

#### 4.5.4 Generalization

To test the models’ capabilities in predicting mutants with a higher number of mutations than they were trained on, the avGFP dataset was used. This dataset is the only one containing variants with up to 14 mutations. Therefore, the training and tune sets consisted of 10,221 and 2,556 data points, respectively, featuring only single and double mutants. The test set consisted of 38,937 variants containing three to 14 mutations. The models were trained under four different settings: from scratch, meaning no pre-training or data augmentation; only with pre-training on our pseudo-score, which contains only scores for single and double mutants; only with data augmentation; and lastly with pre-training and data augmentation combined. The training was done three times with different random seeds to check the prediction consistency. Pearson’s R values between the true and the predicted scores of the test set mutants were computed to evaluate the performance.

#### 4.5.5 Single Mutation Effect Prediction

To test how many training samples a network needs to get an idea of the effect of single mutations, all single mutations of the DMS dataset of GB1 were used as ground truth. Then pre-trained SepConvMixer models were trained on different numbers of training samples of the original datasets (50 -6000 data points). These models, as well as only the pre-trained model of SepConvMixer, were asked to predict the score of every single mutation present. This was done for GB1 because this dataset consists of all possible single mutations, whereas the Pab1 and avGFP datasets would not yield a comparable ground truth due to missing single-point mutations.

#### 4.5.6 Recall Performance

To access the recall performance of simple CNN, SepConvMixer, and DenseNet when trained on different-sized training datasets (Section: 4.5), we used the pre-trained models without data augmentation since this is one of the best-performing settings. The models were trained on different-sized training datasets (50 - 6000 data points) or 80% of the whole datasets. Then the test data set, which consists of only variants that the models have never seen before, was used to access the recall performance by letting the models predict the scores and checking how many of the predicted top-scoring variants were actually part of the actual top scoring 100 variants of the test dataset, given a certain budget (Fig. 6 & S8). The recall performance was computed as described in [18]. If one ranks all variants according to their predicted (fitness-) score, the budget refers to the number of best variants predicted by the network from all variants, which are examined to see whether they occur in the actual 100 best variants. The term “best case” refers to the theoretical optimal outcome. For instance, if we were to select 20 variants from a goal set of the top 100 variants, the best possible outcome would be that all 20 chosen variants are within the true top 100. Thus, the “best case” would reflect a recall score of 0.2. If the budget would be 150 variants, the best case would be that all top 100 variants are contained in the 150 predicted best variants and would therefore result in a recall score of 1.0. The best case is meant as a comparison for what could be the maximum achieved recall score.

## 5 Results

We tested a new way of encoding protein structure and improving the training on deep mutational scanning (DMS) datasets. To this end, a simple convolutional neural network with a LeNet5 [29] -like architecture (simple CNN, Section: 4.3.1), a DenseNet [30] -like Network (DenseNet, Section: 4.3.2) and a network heavily inspired by ConvMixer [31] (SepConvMixer, Section: 4.3.3) were used. Furthermore, two methods, data augmentation and pre-training, were tested for their applicability to DMS data. To assess their performance, a state-of-the-art sequence convolution model [18] (sequence convolution) was trained with the same data sets, and the results were compared. In order to test these models and approaches as well as their real-world applicability, we conducted a series of different experiments to test the following properties:

- the effect of the number of randomly selected training samples as well as of pre-training and data augmentation on the predictive performance (Section: 5.1)
- the ability to extrapolate to unseen sequence positions (Section: 5.2)
- the extent to which the models can predict all single variant effects when trained on an extended alanine scan (ANH-Scan) (Section: 5.3)
- the ability to generalize from training on mutants containing a maximum of two amino acid exchanges to variants carrying up to 14 mutations (Section: 5.5)
- the number of randomly selected data needed to predict the effect of all single mutations (Section: 5.6)
- the recall performance for the best 100 variants in the dataset given a certain budget for networks trained on differently sized datasets (Section: 5.7)

### 5.1 Dataset size effect

Neural networks are known to need a lot of data to perform well. Here we test different network architectures and supplementing methods to reduce the needed data size and its influence on predictive performance. In order to evaluate the performance of the three different architectures and compare it to the original sequence convolution method, each model was trained on three different data sets from [18] (avGFP, Pab1, and GB1), including a limited analysis of the datasets Ube4b and Bgl3. Our study showed improved performance of all three architectures over sequence convolution for smaller datasets and the positive impact of pre-training and data augmentation on their predictive performance. For larger datasets, all models performed almost equally. In order to improve predictions on small data sets, two methods were applied: data augmentation and pre-training. Data augmentation has already been shown to be important when the data set size is small [34]. Since the proposed data representation is not translation and rotation invariant, it was not possible to use the same data augmentation methods (*e*.*g*., rotation, crop, flip, transpose, etc.) as used in image processing. Hence, a simpler data augmentation method was used that sums up scores of existing data (Section: 4.5.1). Another method to improve a model’s performance is pre-training. Here a model can be pre-trained unsupervised if a lot of unlabelled data is available [35] or supervised on a big labeled data set with similar content to the data one is interested in and then fine-tuned on the data set of interest [36]. Since the proposed data representation already captures some variation due to mutations in a protein, we created a simple pre-training procedure, where the models were pre-trained on a pseudo-score that arises from the representations itself (Section 4.5.1).

In the figures 2 and S9 - S13, the median of the three training runs for each data point is shown. The graphs in the top row show either the Pearson-, the Spearman correlation coefficient, or the MSE for the predictions on the test set made by the models. The bottom row shows the relative performance compared to the sequence convolution. The data set size always refers to the amount of data from the original split and is not related to the data set size after data augmentation. For the MSE, the relative performance was calculated with *p*_MSE_ = 2 ***−*** (*MSE*_i_*/MSE*_seqconv_) where *p*_MSE_ is the relative performance, the *MSE* _i_ the MSE of a model to compare to and *MSE* _seqconv_ the MSE of the sequence convolution. The relative performance of the correlation coefficients was calculated *p*_R_ = *R*_i_*/R*_seqconv_ where *p*_R_ is the relative performance, *R*_*i*_ the correlation coefficient for the model to compare to and *R*_*seqconv*_ the correlation coefficient of the sequence convolution model.

**Figure 2:**
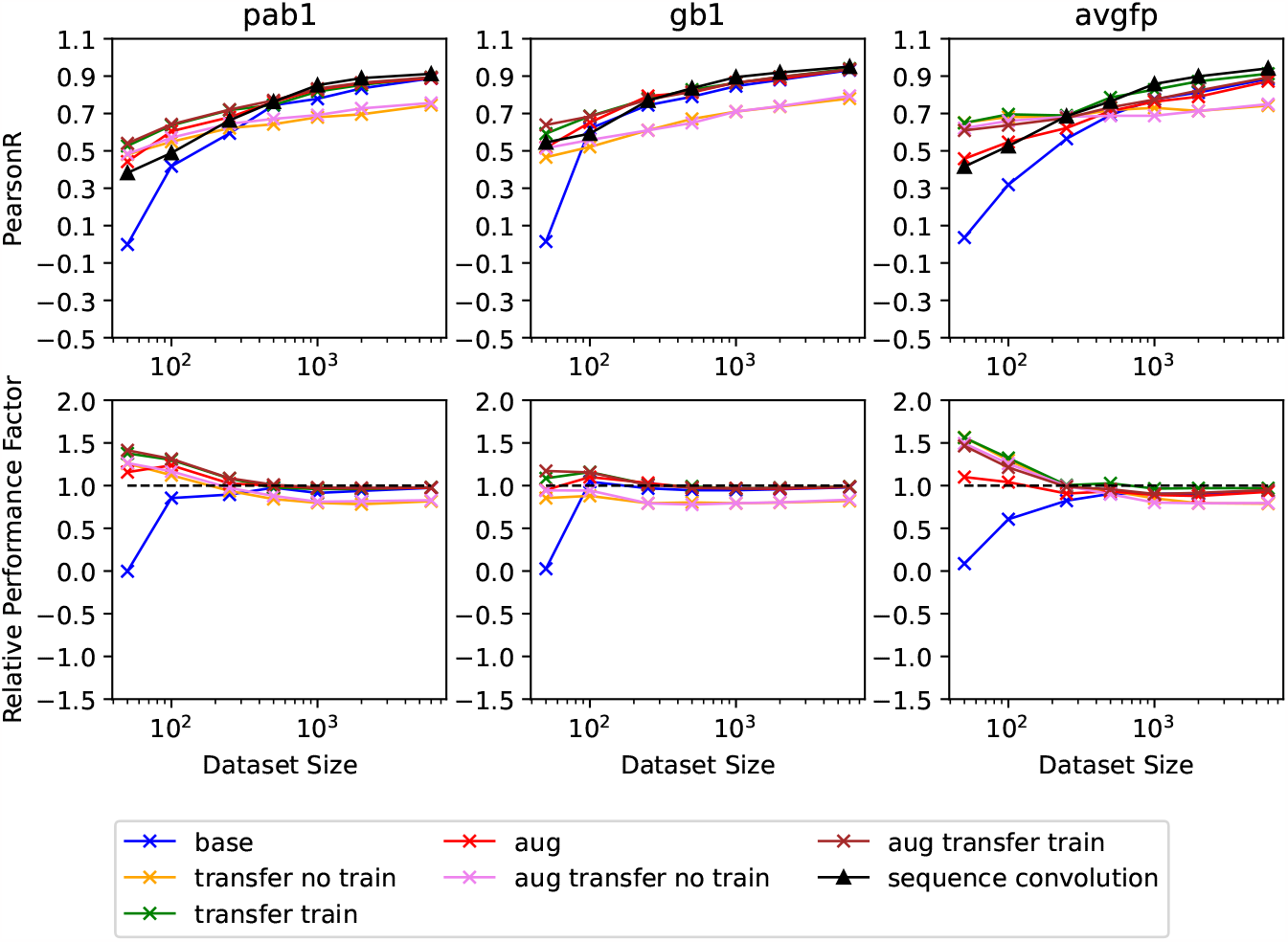
Pearson’s R for predictions of the test data set of SepConvMixer for all three proteins in the upper row, as well as the relative performance compared to sequence convolution in the lower row. Here sequence convolution is indicated as a black dashed line at 1×of its own performance. Label descriptions can be found in Table 3

Augmentation specifies whether data augmentation was used, transfer whether pre-training was used, and train CL whether the convolution layers were trainable or not when pre-training was used.

**Table 3:**
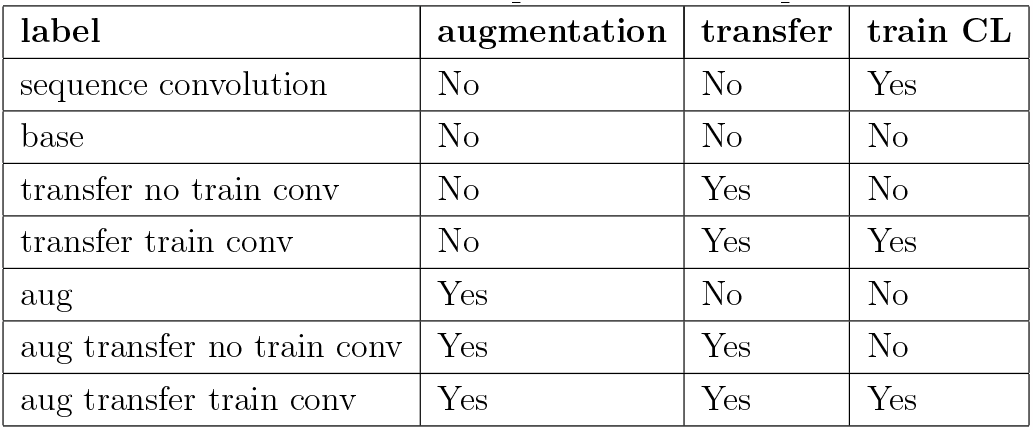
Label description for result plots.

| label | augmentation | transfer | train CL |
| --- | --- | --- | --- |
| sequence convolution | No | No | Yes |
| base | No | No | No |
| transfer no train conv | No | Yes | No |
| transfer train conv | No | Yes | Yes |
| aug | Yes | No | No |
| aug transfer no train conv | Yes | Yes | No |
| aug transfer train conv | Yes | Yes | Yes |

In general, the more data the networks got to train, the better they performed, and the less important the approach became since they performed almost equivalently (*>*= 2000 training samples). Another trend that could be observed is that the more original data the networks got, the less important augmentation and pre-training became to achieve the same training results. In general, the best performances were obtained when the networks were pre-trained, and the weights of the convolutional layer were not frozen in the subsequent training runs. Data augmentation had an additional positive effect. On smaller datasets (*<*= 500 training samples), the difference in the performance of a chosen method was more pronounced. For example, for Pab1 and avGFP, using data augmentation and freezing the convolutional layers showed a better performance in simple CNN (Fig. S15) but showed a worse performance when the training dataset got bigger. In contrast, this method led to an overall worse performance in DenseNet (Fig. S16) and SepConvMixer. This was especially true for SeqConvMixer and could be caused by the low number of trainable parameters (13k) for the network under this setting (Fig. 2). Looking at the method that produced the best results, training a pre-trained network and using data augmentation, DenseNet had a similar performance overall to simple CNN and SepConvMixer. A performance improvement from DenseNet could be seen in small datasets (Fig. S17).

When the number of training samples gets over 500, the performances of all architectures are almost identical. One fact that stood out about DenseNet was that it took at least 6000 samples to show the same performance as sequence convolution when no pre-training and data augmentation were used. In contrast, simple CNN without pre-training and data augmentations needed 250 to 500 training samples to show the same performance as sequence convolution. In general, the difference in performance for different methods was less pronounced in simple CNN than in DenseNet and SepConvMixer. Looking at the difference in performance between simple CNN, SepConvMixer, and DenseNet, one can see that DenseNet could improve the performance for smaller datasets when pre-training and/or data augmentation was used. On the other hand, when none of these methods were used, DenseNet showed a strongly reduced performance and higher variability in its results (Figs. S19 - S21). Data augmentation worked well for data sets that feature only single and double mutants, such as Pab1 and GB1. When the dataset already consisted of variants with more than two mutations (up to 14 in the case of avGFP), the approach did not work as well when the training data size surpassed 250 entries. This might be caused by the additive nature of the data augmentation used in our pipeline. Since adding two single mutants is more likely to be additive in real life compared to adding two variants, both carrying 12 mutations on their own, because a higher number of variants increases the likelihood that two mutations interfere with each other and, therefore, corrupt the additivity of their scores when they occur on their own.

For the lower-quality datasets of Ube4b and Bgl3, we performed a limited analysis with only our biggest and smallest architecture, simple CNN and SepConvMixer, and only two training settings, without pre-training and data augmentation, and with pre-training. We could see an increase in performance when pre-training was used, but as already shown in [18], we could observe the same trend with a reduced performance compared to the other three datasets (Fig: S18).

Number of trainable parameters of the three different architectures for two different proteins: Pab1 (75 amino acids) and avGFP (237 amino acids).

### 5.2 Positional Exploration

Since training on a randomly chosen subset of data points can be biased by the fact that it already learned that a mutation at a particular position will produce a bad result, we trained our smallest and biggest network architecture on data of different sequence positions than they were asked to predict (Figure 3 and Figure S23). Even though the networks were trained on bigger datasets than in Section 5.1 (23000 Pab1, 26000 avGFP, and up to 351000 for GB1), they showed a worse performance compared to substantially smaller training data that did not exclude specific positions. Here, simple CNN and SepConvMixer show comparable performance. Both approaches manage to improve over the predictions made when trained on the protein sequences with sequence convolution.

**Figure 3:**
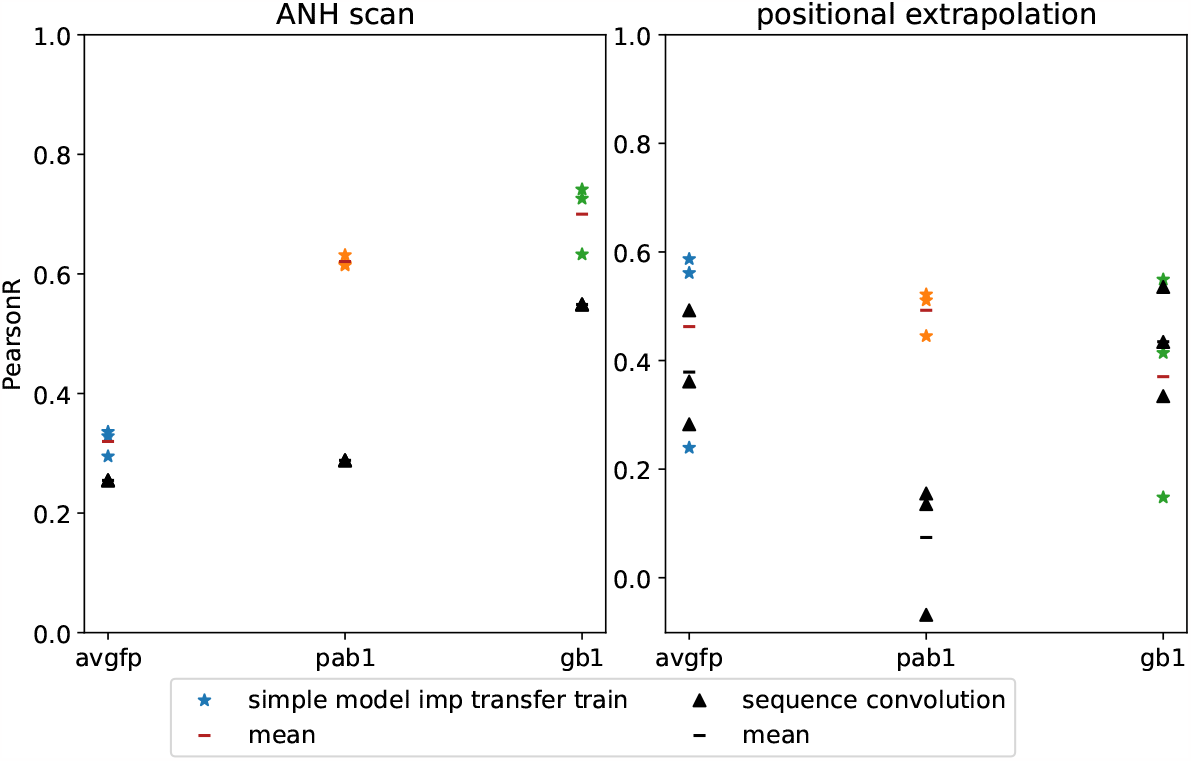
PearsonR for predictions on an ANH-Scan as well as on positions the networks (simple CNN and sequence convolution) have not seen before in training

Since the size of the training dataset was big enough, there were no big differences in performance between pre-trained and not pre-trained networks(Fig. S23 & S25).

### 5.3 ANH-Scan

Performing an alanine scan on a protein will only yield a small number of data points. Therefore, an extension to systematically replace each amino acid with alanine, as well as asparagine and histidine, was tested to see how well the networks could predict individual amino acid replacements with the remaining amino acids (Figure 3 and Figure S23). We tested this approach with simple CNN, SepConvMixer, as well as with sequence convolution. This approach showed similar results to the randomly chosen single- and multi-mutation variants in Section 5.1 for simple CNN and SepConvMixer and a slight performance degradation for sequence convolution for similar-sized train datasets. Here the reduced performance on not pre-trained networks, again, shows its importance when training data is limited.

### 5.4 Comparison - Sequence vs Structure

In general, comparing the correlations of all three networks to sequence convolution for predictions when trained on randomly chosen positions (Section 5.1), the best combination of methods (using pre-training with and without data augmentation) performed at worst 0.9×of sequence convolution and the best 1.6×. Using a more complex architecture (DenseNet) could improve the performance on smaller dataset sizes (*<*= 250) but needed at least pre-training to reach that level of performance (Fig. S17). When comparing the number of parameters (Table: 4) for sequence convolution and simple CNN, the protein sequence length is the determining factor. The bigger the protein, the more will the simple CNN exceed the sequence convolution in terms of the number of parameters. For DenseNet and SepConvMixer, the number of parameters stayed constant and only changed due to the use or absence of the first introduction layer. This difference was due to the use of a flatten layer in the sequence convolution and simple CNN after their feature extraction part, whereas DenseNet and SepConvMixer both use a global pooling layer instead that always has the same size, regardless of the input data.

**Table 4:** Number of parameters of each network.

| architecture | protein | trainable parameter |
| --- | --- | --- |
| sequence convolution | Pab1 | 990k |
| simple CNN | Pab1 | 803k |
| DenseNet | Pab1 | 714k |
| SepConvMixer | Pab1 | 37k |
| sequence convolution | avGFP | 3.118k |
| simple CNN | avGFP | 7.029k |
| DenseNet | avGFP | 799k |
| SepConvMixer | avGFP | 38k |

Comparing the ability to correctly predict the effect of amino acid exchanges at positions that were not present in the training data, sequence convolution, as well as our models, decrease in performance, especially considering the overall larger training data set. Our combination of structure encoding and pre-training managed to slightly improve predictions on avGFP, improve predictions on Pab1, and perform slightly worse on GB1 in comparison. Simple CNN managed to perform the same or with a 0.45 higher PearsonR compared to sequence convolution without pre-training. SepConvMixer achieves, at worst, a 0.12 lower PearsonR or, at best, a 0.49 higher PearsonR (Fig. S24 & S25)

When the training data came from a simulated ANH scan, sequence convolution lagged behind both of our models in terms of predicting the single mutation effect of the remaining amino acids when they were pre-trained but outperformed them when they were not pre-trained.

### 5.5 Generalization

Examining the performance of models trained on only single and double mutants in predicting mutants that have more than two amino acid exchanges again showed the advantage of pre-training and data augmentation. The various networks (simple CNN, DenseNet, SepConvMixer) were trained on 10.221 single and double mutants of avGFP, with and without pre-training and/or data augmentation. Then they were asked to predict the test dataset that contains variants featuring a minimum of three and a maximum of 14 mutations. This led to a maximum performance in terms of Pearson’s R of 0.835 (Fig. 4). Over all settings and methods, simple CNN showed the best results, followed by SepConvMixer, whereas DenseNet showed the worst performance. Looking at the consistency of the results, SepConvMixer outperforms the other two networks. In accordance with previous results, the models can improve their predictions when pre-training and data augmentation are used. Also in line with our previous experiments, the best setting combination was pre-training combined with data augmentation.

**Figure 4:**
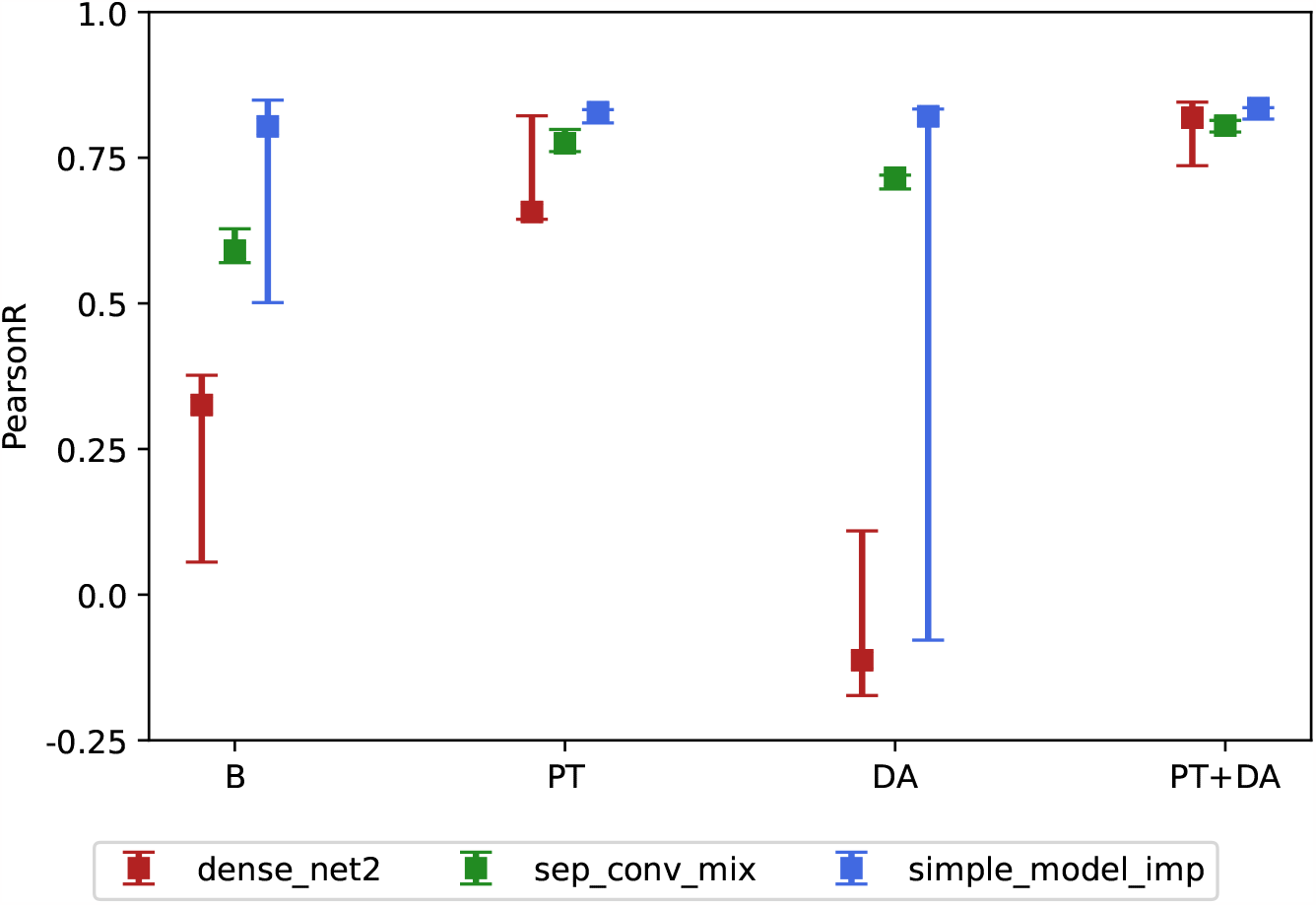
Pearson’s R of predictions for variants of avGFP containing three to 14 mutations when the networks were only trained on single and double mutants (B: no pre-training and no data augmentation, PT: with pre-training, DA: with data augmentation, PT+DA: with pre-training and data augmentation)

Under these settings, all models performed the best and delivered the same performance.

### 5.6 Single Mutation Effect Prediction

Testing the performance on predicting the effects of single mutations of pre-trained SepConvMixer networks trained on reduced dataset sizes (Section: 4.5) showed under visual comparison that for GB1, a protein with a sequence length of 56 amino acids, models trained on 250 training samples started to have a good idea of which single mutations had a positive and which had a negative effect (Fig. 5). This comparison was only possible for GB1 since its data set contained possible single mutants.

**Figure 5:**
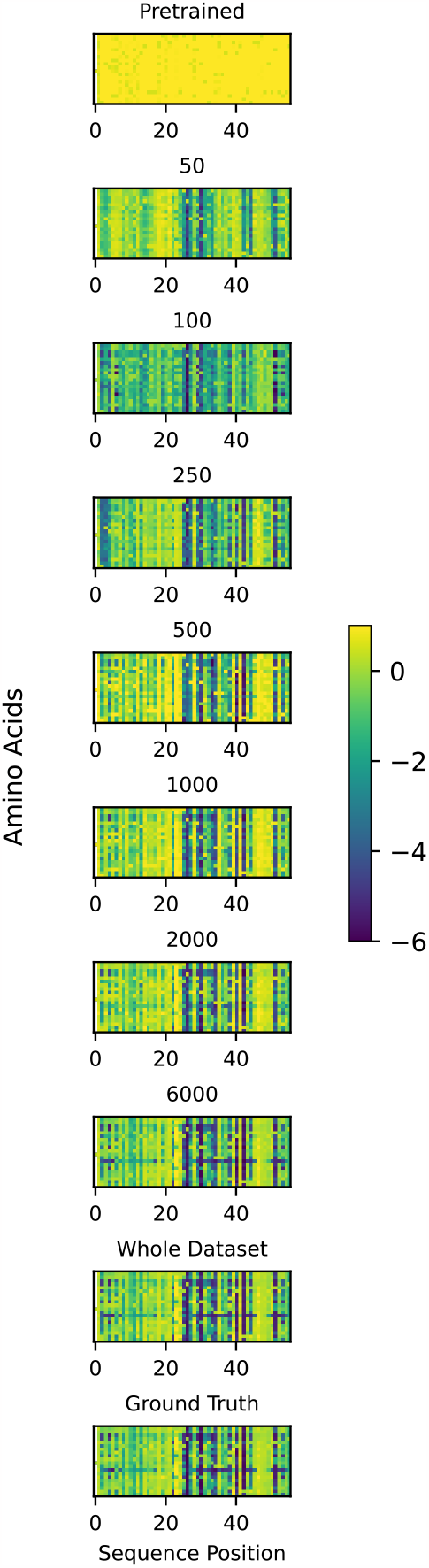
Prediction of the mutational effect of each single mutation at each sequence position using SepConvMixer on the example of GB1. The pre-trained models were trained on training data sets consisting of 50 to 6000 or 80% of the whole GB1 dataset and asked to predict the score of every possible single mutation of GB1. For comparison, the actual measured data are shown as ground truth and the result of a model that was only pre-trained. Figure S22 shows the difference of all predictions to the ground truth. On the y-axis, the amino acids are alphabetically ordered.

### 5.7 Recall Performance

Asking networks to recall the top 100 variants given a certain budget showed some differences for the different datasets but smaller differences in performance between all network architectures (Fig. S8). The recall performance on Pab1 and avGFP showed similar results, whereas the recall for GB1 showed better recall results when trained with the same train dataset size. The overall trend showed that it is advantageous to invest in more training data to then be able to better recall the true top variants. For Pab1 and avGFP, a bigger increase in performance could be seen when changing from 6000 training samples to the whole (80% of the whole data set) dataset, whereas, for GB1, no performance increase between 6000 training samples and the whole dataset could be seen. Comparing the results for when trained with different amounts of training data, SepConvMix trained on 6000 training samples, needed a budget size between 70 to 1040 samples to recall 60 % of the top 100 variants. When trained on 500 training samples, it needed between 500 to 1270 samples to match this performance, and when trained on 50 training samples, 1390 up to 1700 samples were required to reach this performance (Fig. 6).

**Figure 6:**
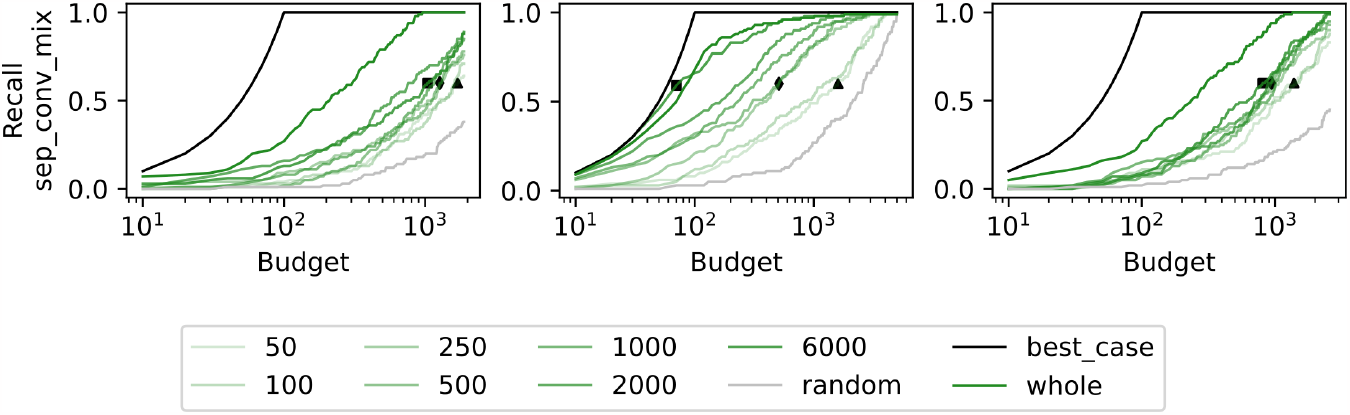
Recall of the top 100 test set mutations given a certain budget (number of predictions that may contain the true Top 100) for SepConvMixer. The models were trained on train data sets containing 50 - 6000 data points or on 80% of the whole data set, which is labeled “whole”. The 60% recall performances when trained on 6000 data points are shown as ◼, as ♦ when trained on 500 data points, and as ▲ when trained on 50 data points. The term “best case” refers to the theoretical optimal outcome. (For a detailed description, see Section 4.5.6)

## 6 Discussion

It has previously been shown that incorporating the structure in the form of a graph and training a graph neural network to predict deep mutational scanning results achieves the same performance as sequence convolution [18]. By introducing our new protein structure representation, we could show that it contains valuable information, can create pre-training data without any experimental data needed, and can improve predictive performance when positions were not seen during training. We could show that the predictive performance could be greatly improved by two straightforward but effective methods, pre-training and data augmentation. Even though the pre-training is very effective, it is not enough to freeze the convolution layers and only let the fully connected layers be trainable. This showed already worse performance after the train sample size exceeded 500 samples, even though simple CNN was able to compensate better because the major part of its architecture consists of fully connected layers (Section: 5.1).

When deciding how to generate a dataset for the optimal outcome of training a neural network, the comparison between ANH-Scan and positional extrapolation (Section: 5.2 & 5.3) showed that a comparable result to a big dataset (positional extrapolation) can be achieved with only a fraction of the data needed (ANH-Scan) when all positions are present in the training data. Even better results can be achieved, also with a fraction of the dataset needed when randomly chosen single- and multi-mutational variants are used (Section: 5.1). By testing the recall performance, it became clear that it is advantageous to invest in more training data because the networks will then be better at predicting the true best variants (Section 5.7).

Data augmentation can be advantageous when used with datasets containing only single and double mutants and network architectures with a smaller number of parameters. When the network architecture with the highest number of parameters was used (Simple CNN for avGFP), one can see that this network, when no pre-training is used, over-fitted the augmented data. Since this data resembles synthetically generated fitness scores that are not always correct, it has to be used with architectures that use fewer parameters. In contrast, the smallest network (SepConvMixer) still managed to perform decently when only data augmentation was used (Section 5.5).

When comparing the performances on predictions on unseen positions, one can see that our encoding either performs the same without pre-training or shows improved performance over a sequence input, suggesting that the encoding enables better extrapolation due to the encoded interactions between amino acids.

Interestingly, SepConvMixer performed almost the same as the other architectures despite its much smaller number of parameters. This is promising since fewer parameters reduce the risk of over-fitting. Therefore, the network should be better able to generalize to unseen data. Furthermore, this network will need fewer computational resources. We also showed that more modern network architectures compared to simple CNN, could improve the performance when the training sample size is small. Since no dedicated hyper-parameter tuning was performed, an increase in the performance of the models is still possible.

The current way our contact maps are generated is used as a fast and simple approximation of the changes happening in the structure of a protein due to amino acid substitutions. The matrix representing the charge interactions does not take into account the protonation state of the amino acids, and the hydrophobicity matrix is only a simple scale and does not take the side chain surroundings into account. These are just two examples where improvements in the protein structure representation can still be made. This might, in turn, help the network to predict mutational effects even better due to a more realistic representation. Using different protein sequence alignment databases (non-redundant and experimental, which is a 90% clustered version of the non-redundant database) did not change the training results significantly. An advantage of our encoding is the possibility to encode “average” structures derived from molecular dynamics simulations or structures with optimized rotamer positions which leads to an even more natural representation of the protein structure and could potentially create an even better encoding through the interaction matrices.

With the advancement of programs like AlphaFold [23] and RoseTTAFold [37], we can assume that there is a trustworthy structure for most proteins. Even homology modeling might be sufficient to supply a decent protein structure that can be used to create the structure representation.

Besides the advantage of being able to use a large number of different architectures derived from the computer vision field, our encoding has the additional advantage of being computationally efficient while representing the biophysical-, interaction- and structural change that occurs due to amino acid substitutions. This makes a more structure-related workflow feasible for researchers without access to high-performance (computing) clusters.

Regarding experimental protein engineering in the lab, minimizing the data size required to achieve comparable or superior prediction results is crucial in reducing time, cost, and resources. Our analysis indicates that these models already perform reasonably well in that respect. They could also be utilized for datasets that do not originate from DMS but rather from a conventional “low throughput” experiment like ANH-Scans, thus providing well-trained mutation effect oracles to more laboratories.

The ultimate goal would be to transfer “learnings” from one DMS dataset to train a universal network capable of predicting the fitness of proteins for which no experimental data exists is very intriguing but currently most likely restricted to proteins with sufficiently similar structures.

## Conflict of Interest Statement

The authors declare that the research was conducted in the absence of any commercial or financial relationships that could be construed as a potential conflict of interest.

## Author Contributions

GW: Conceptualization, Data curation, Investigation, Methodology, Software, Validation, Visualization, Writing – original draft, Writing – review & editing, Project administration IP: Conceptualization, Methodology, Writing – original draft GO: Conceptualization, Methodology, Writing – original draft, Writing – review & editing KG: Conceptualization, Methodology, Writing – original draft, Writing – review & editing, Funding acquisition, Project administration, Resources, Supervision, Validation

## Funding

Funding was provided by the Austrian Science Fund (FWF) through project DOC-130 (doc.funds BioMolStruct - Biomolecular Structures and Interactions) and by the Doctoral Academy Graz of the University of Graz.

## Acknowledgement

The authors thank Prof. Thomas Pock and Prof. Robert Kourist for fruitful discussions.

## Data Availability

The data used to train all networks and the code can be found on GitHub (https://github.com/ugSUBMARINE/image-dms) and is licensed under the MIT license. The code is set up so that it can easily be used to reproduce the results of this publication. It can also be easily adapted to pre-train and train our networks (or any other network of interest) on a new dataset.

## Notes

https://github.com/ugSUBMARINE/image-dms

### Competing Interest Statement

The authors have declared no competing interest.

### Summary of Updates

Several text and format changes.

