## Supporting Information (25 figures) for "Flattening the curve - How to get better results with small deep-mutational-scanning datasets"

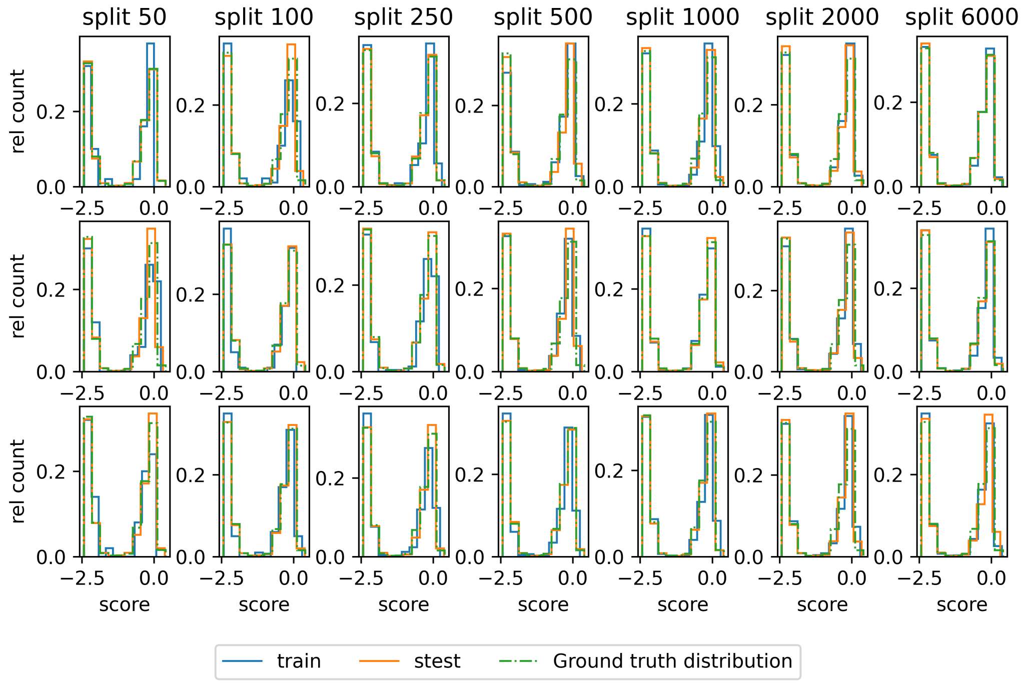


Figure S1: DMS score distribution of the training- and test dataset as well as the distribution of the whole dataset (ground truth distribution) for avGFP for one of the training triplets. The counts were divided by the dataset size to be able to compare the distributions

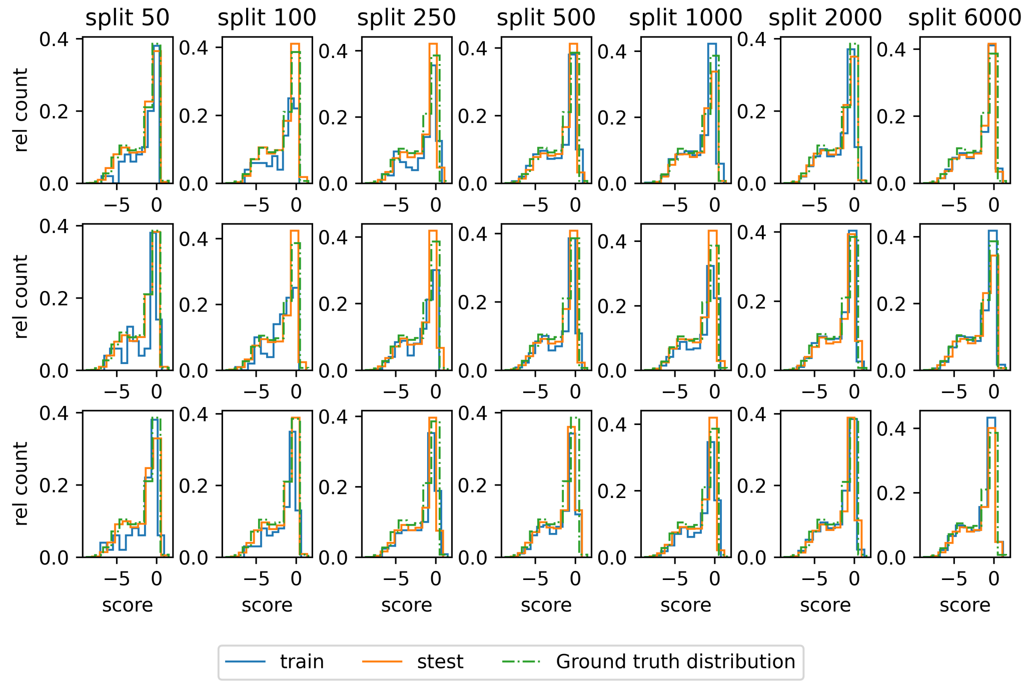


Figure S2: DMS score distribution of the training- and test dataset as well as the distribution of the whole dataset (ground truth distribution) for Pab1 for one of the training triplets. The counts were divided by the dataset size to be able to compare the distributions

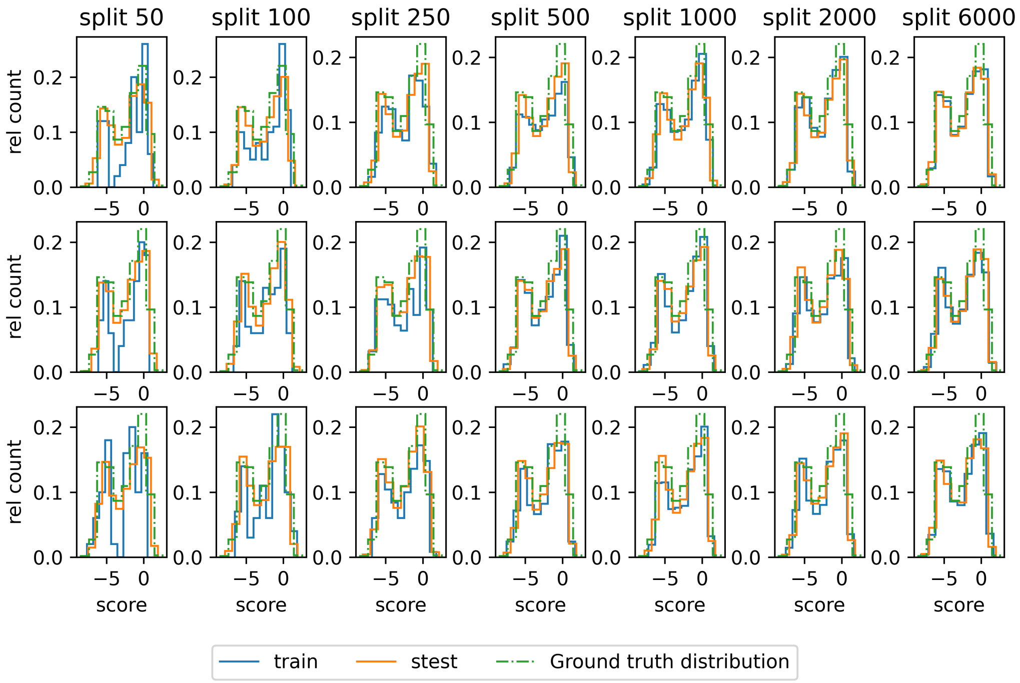


Figure S3: DMS score distribution of the training- and test dataset as well as the distribution of the whole dataset (ground truth distribution) for Gb1 for one of the training triplets. The counts were divided by the dataset size to be able to compare the distributions

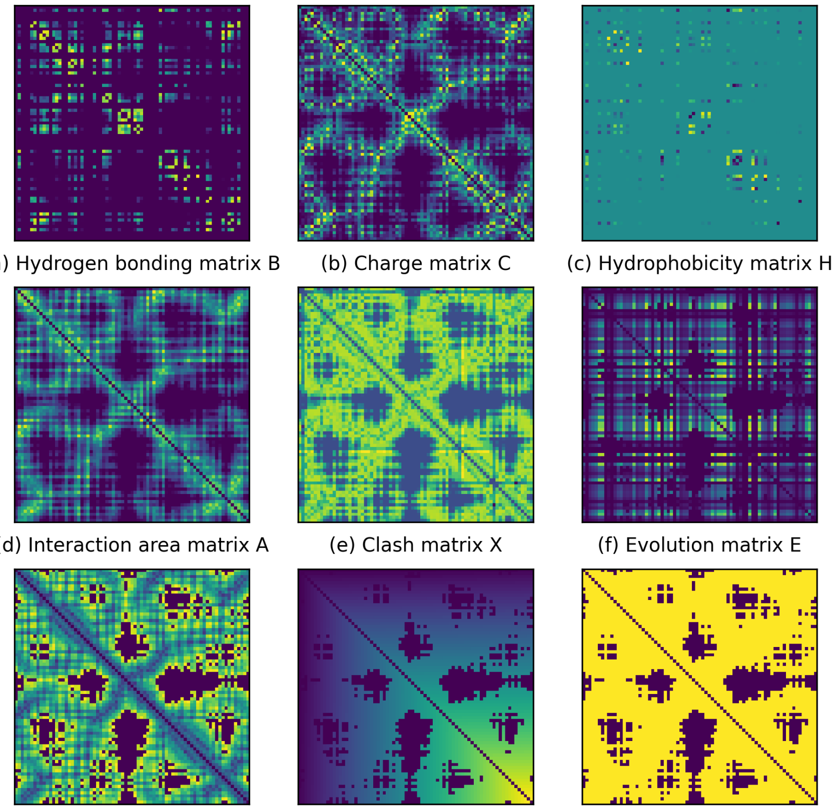

(a) Hydrogen bonding matrix B (b) Charge matrix C (c) Hydrophobicity matrix H

(d) Interaction area matrix A (e) Clash matrix X (f) Evolution matrix E

(g) Distance matrix D (h) Position matrix P (i) Interaction Matrix M

Figure S4: Plots of all interaction describing matrices as well as distance, the position and the interaction (distance matrix with applyed $d_{th}$) matrices of Pab1 containing the mutations "N127R, A178H, G177S, A178G, G188H, E195K, L133M, P125S"

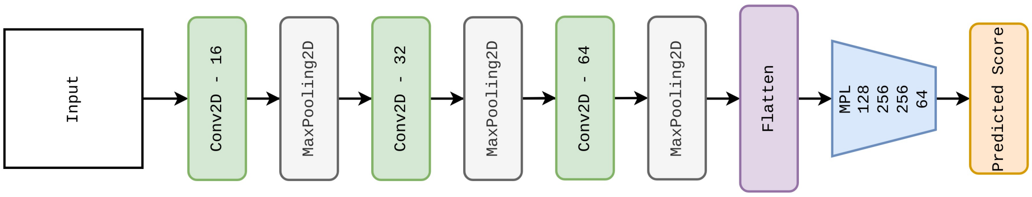


Figure S5: SimpleCNN architecture

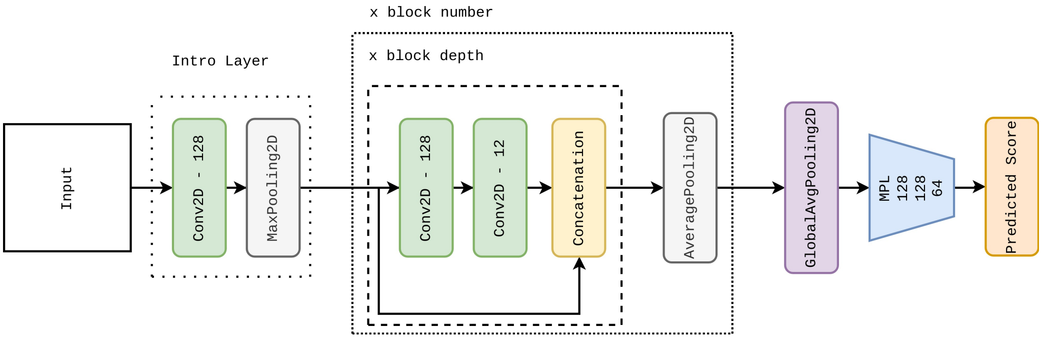


Figure S6: DenseNet architecture

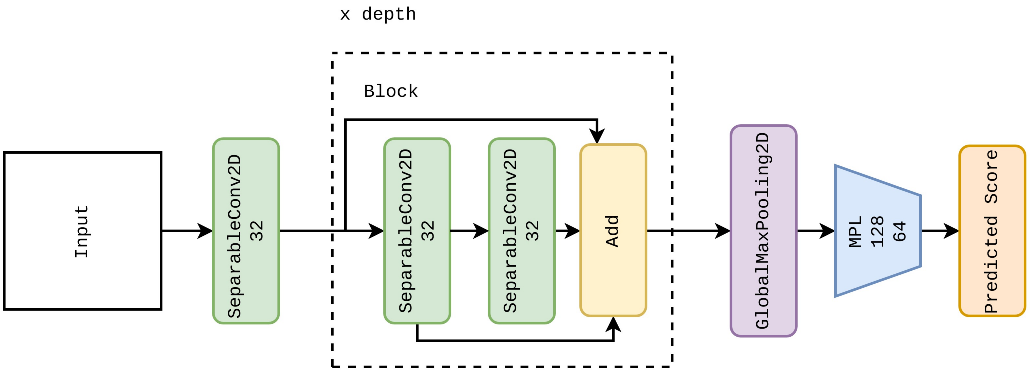


Figure S7: SepConvMix architecture

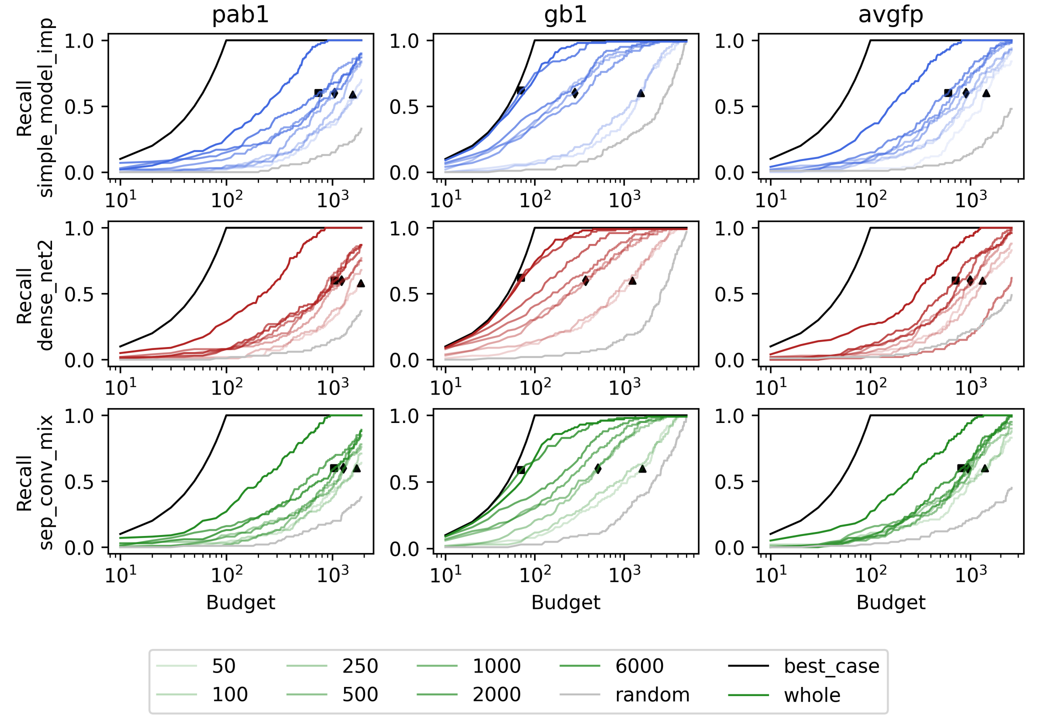


Figure S8: Recall of the top 100 test set mutations given a certain budget (number of predictions that may contain the true Top 100) for SepConvMixer. The models were trained on train data sets containing 50 - 6000 data points or on 80% of the whole data set, which is labelled ”whole”. The 60% recall performances when trained on 6000 data points are shown as ■, as ⧫ when trained on 500 data points and as ▲ when trained on 50 data points.

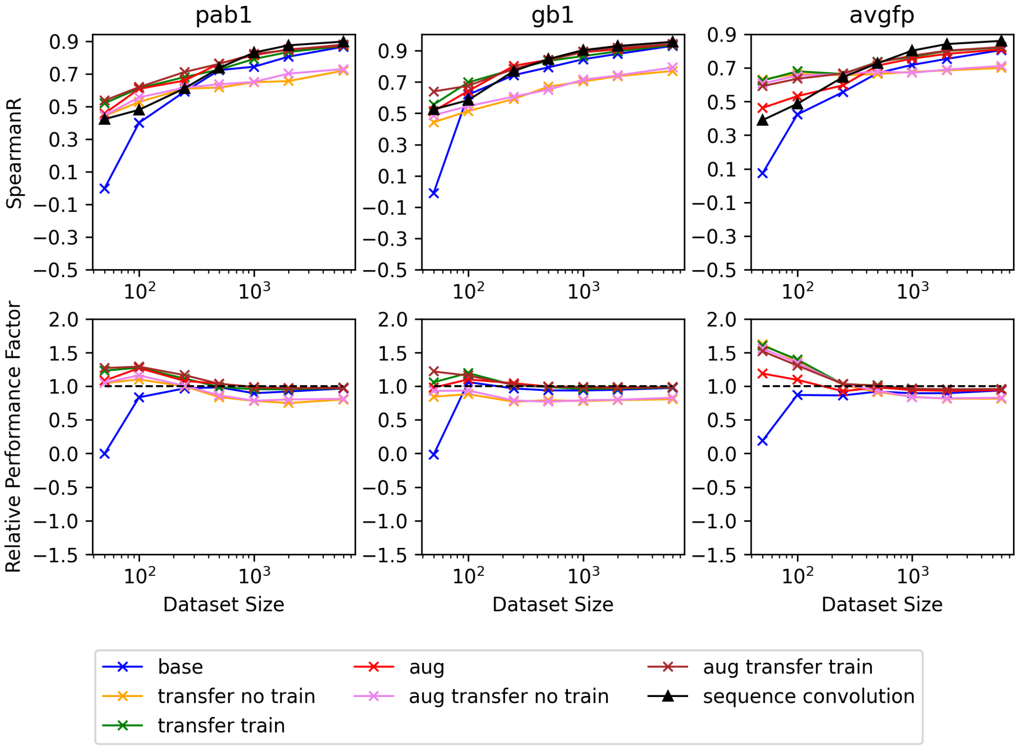


Figure S9: Spearman R for predictions of the test data set of SepConvMixer for all three proteins

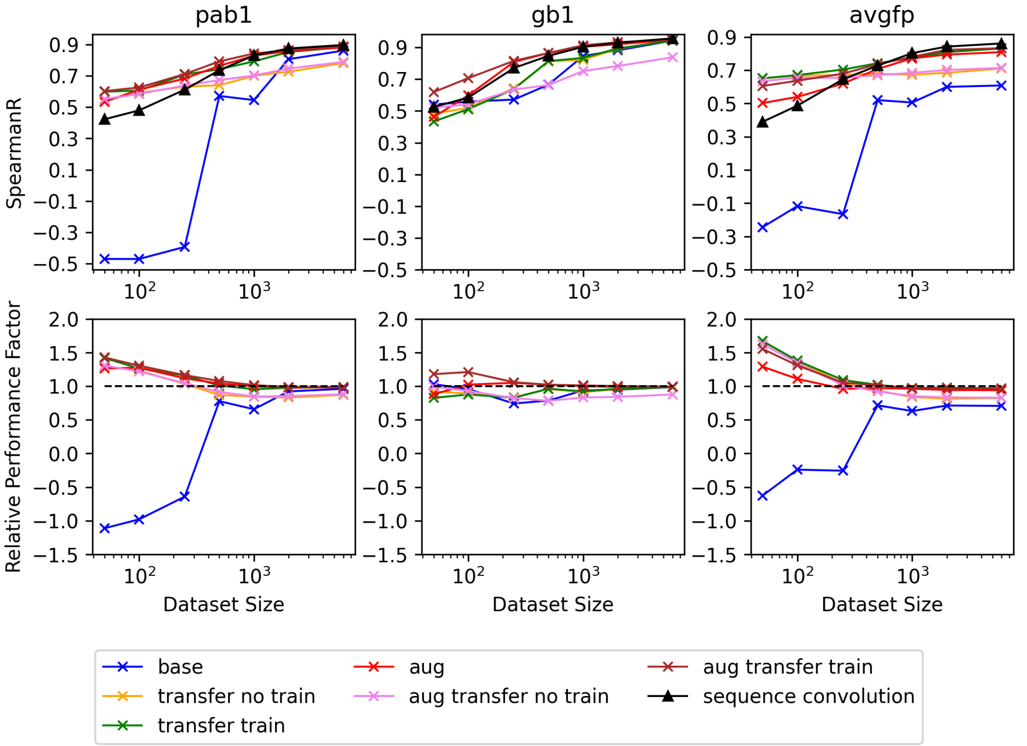


Figure S10: Spearman R for predictions of the test data set of DenseNet for all three proteins

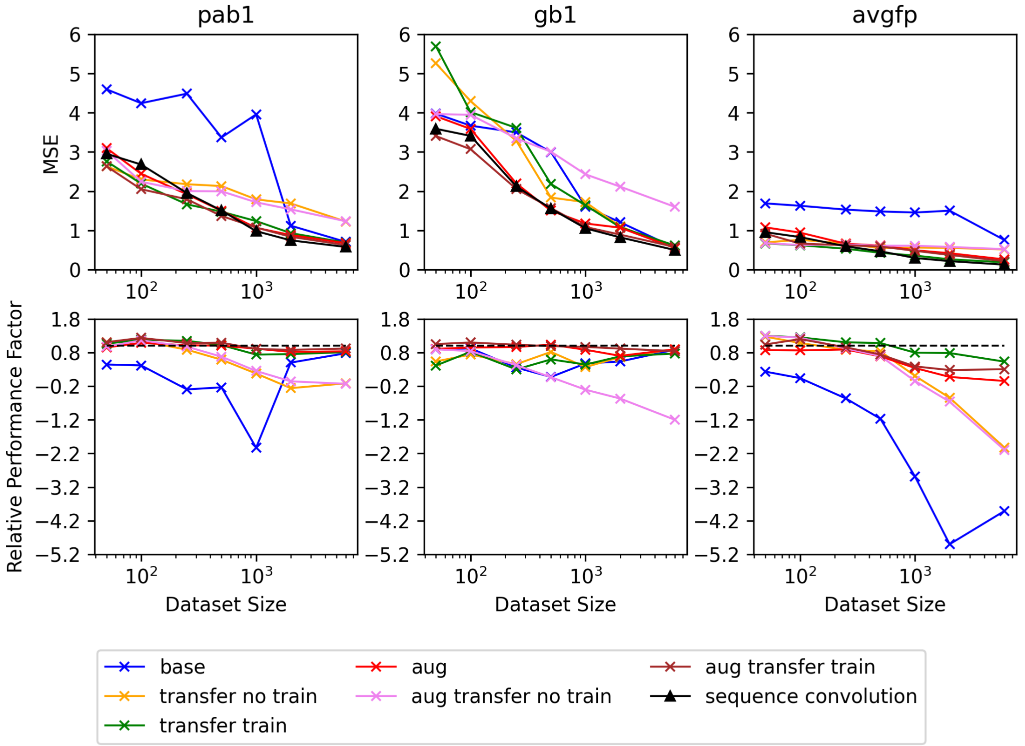


Figure S11: MSE for predictions of the test data set of DenseNet for all three proteins

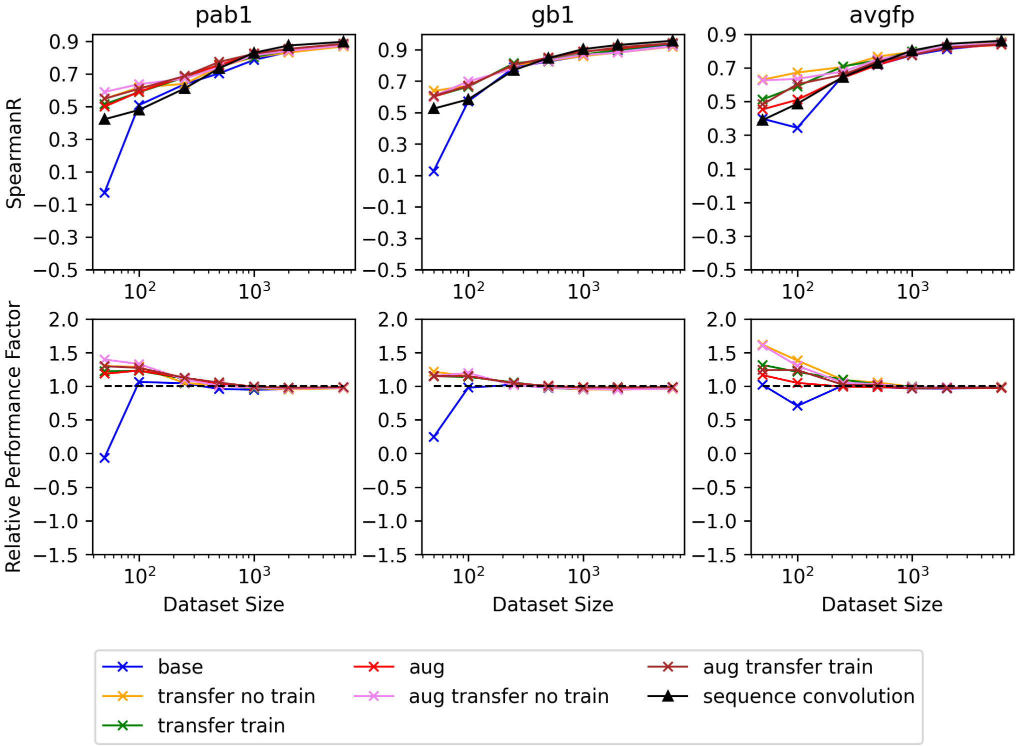


Figure S12: Spearman R for predictions of the test data set of simple CNN for all three proteins

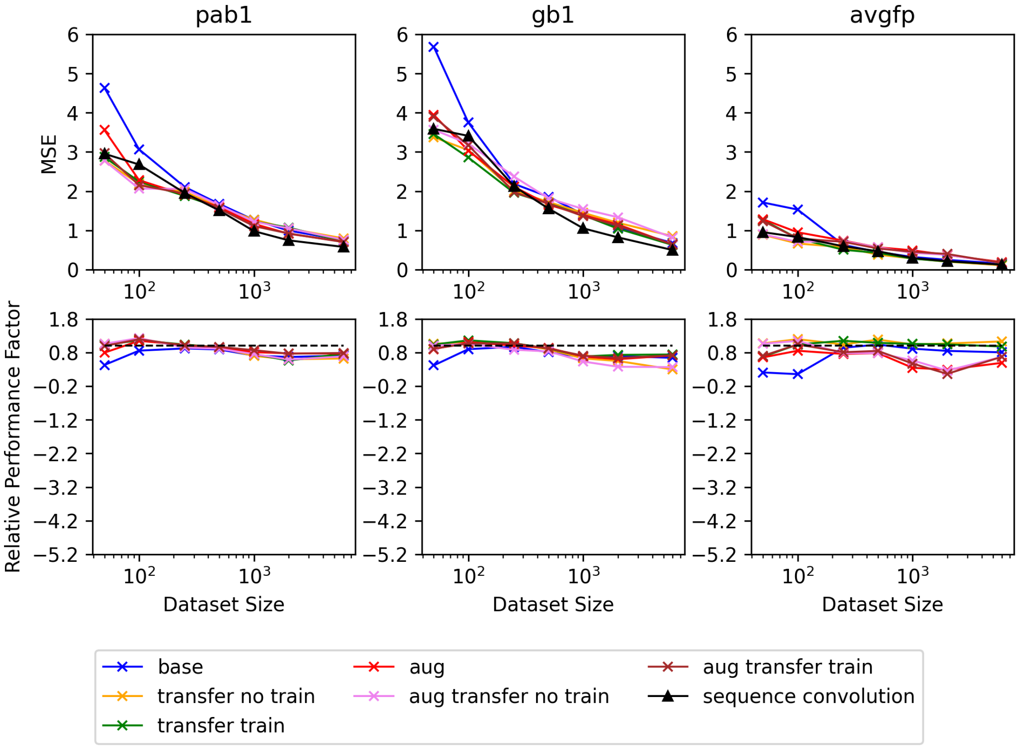


Figure S13: MSE for predictions of the test data set of simple CNN for all three proteins

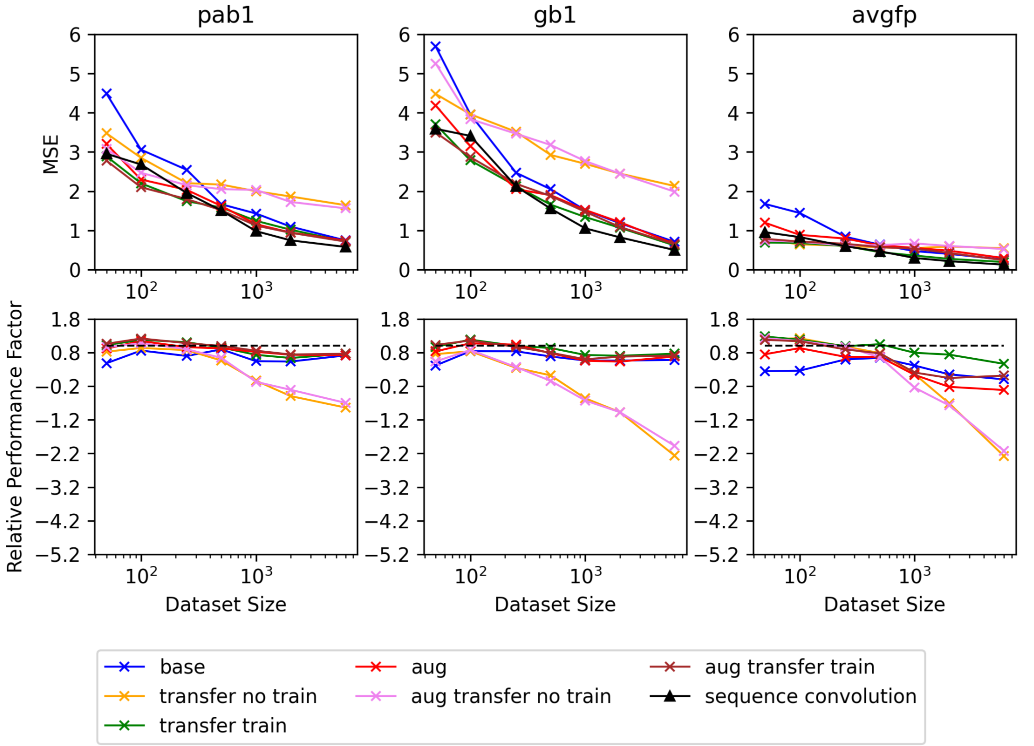


Figure S14: MSE for predictions of the test data set of SepConvMixer for all three proteins

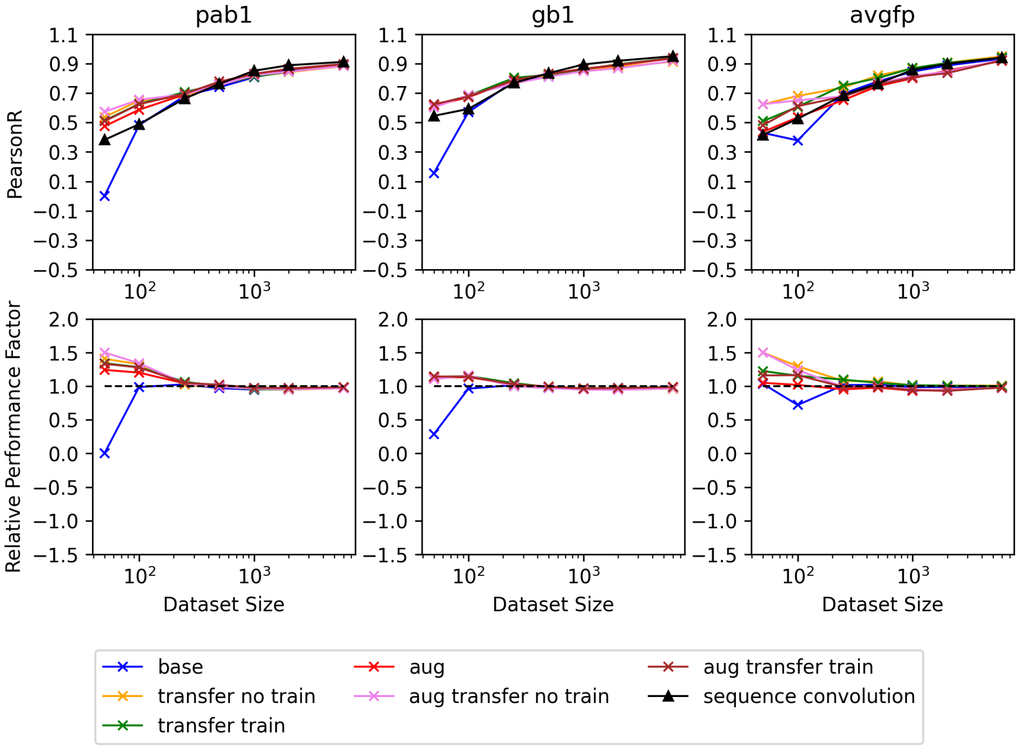


Figure S15: Spearman R for predictions of the test data set of simple CNN for all three proteins

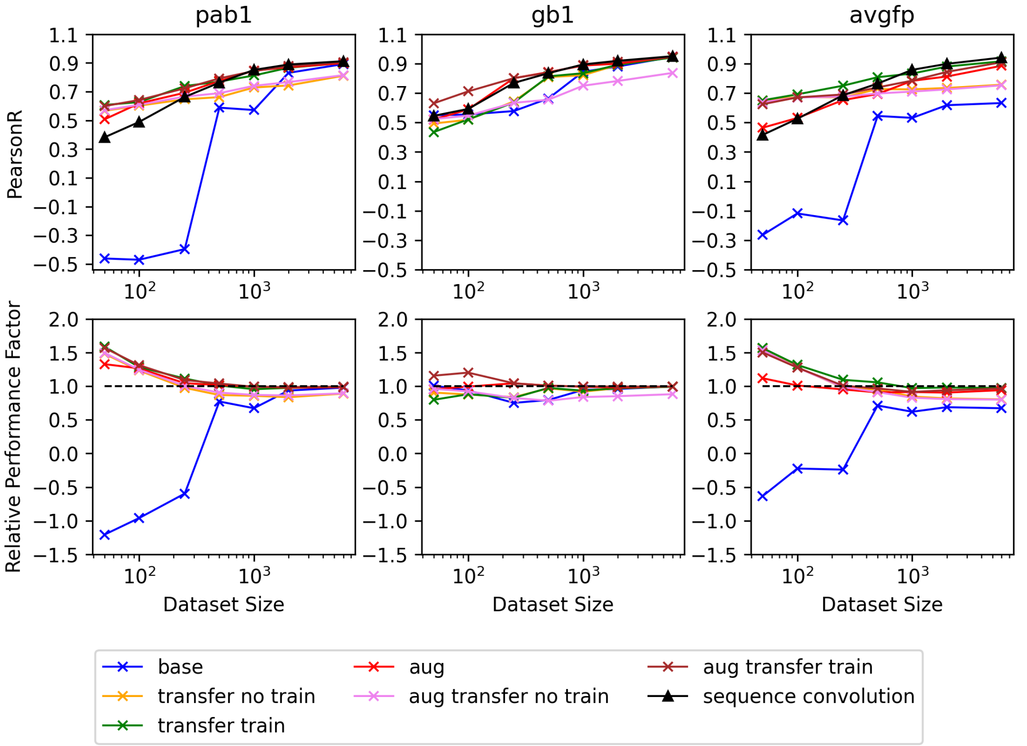


Figure S16: Spearman R for predictions of the test data set of DenseNet for all three proteins

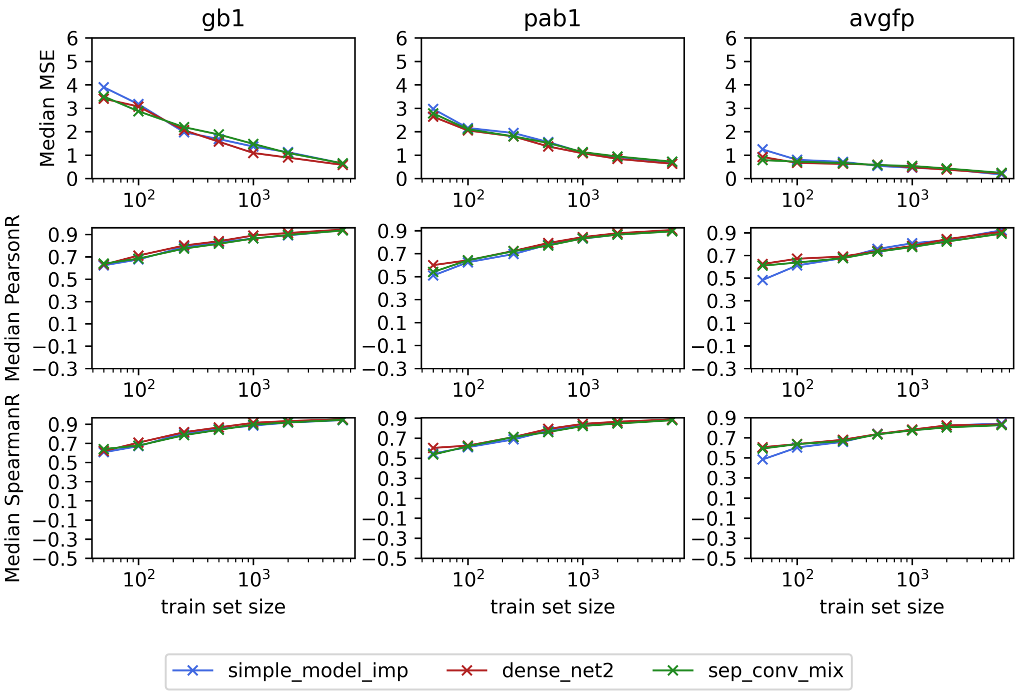


Figure S17: Comparison of training results for "aug transfer train conv" for all three datasets between simple CNN and DenseNet

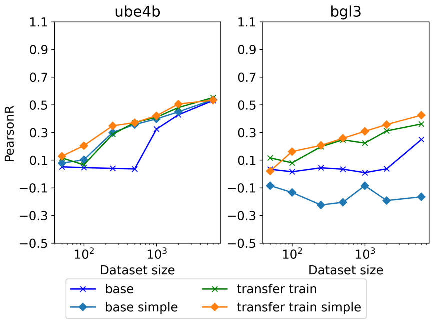


Figure S18: Pearson R for predictions of the test data set of Ube4b and Bgl3 using SepConvMixer without pretraining and without data augmentation (base) and pretrained on the pseudo-score with a trainable feature extractor during the training on the DMS data (transfer train)

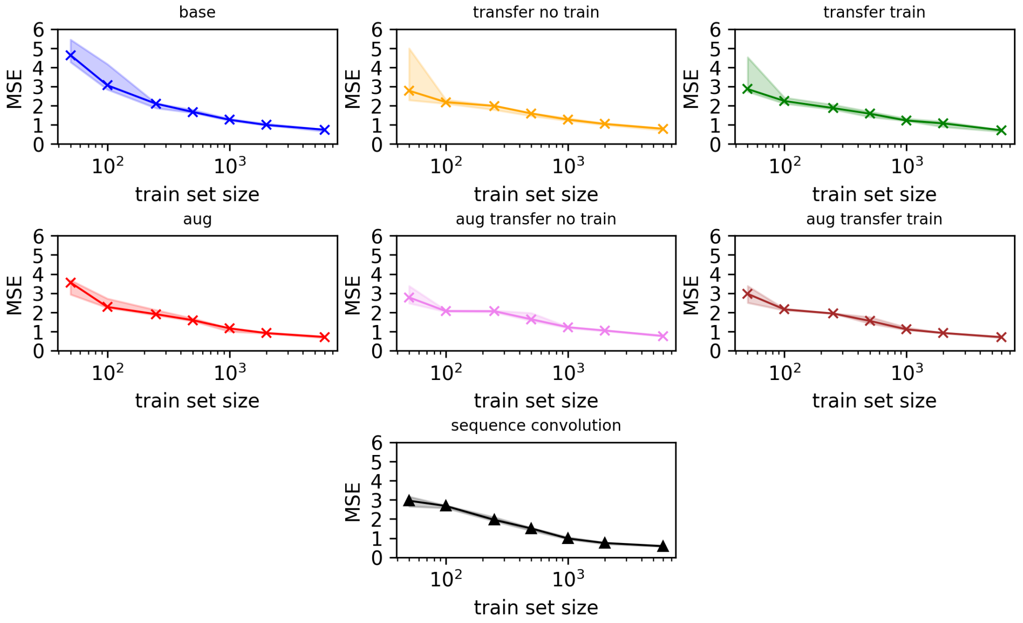


Figure S19: Spread of MSE values of the different training runs for simple CNN trained on Pab1

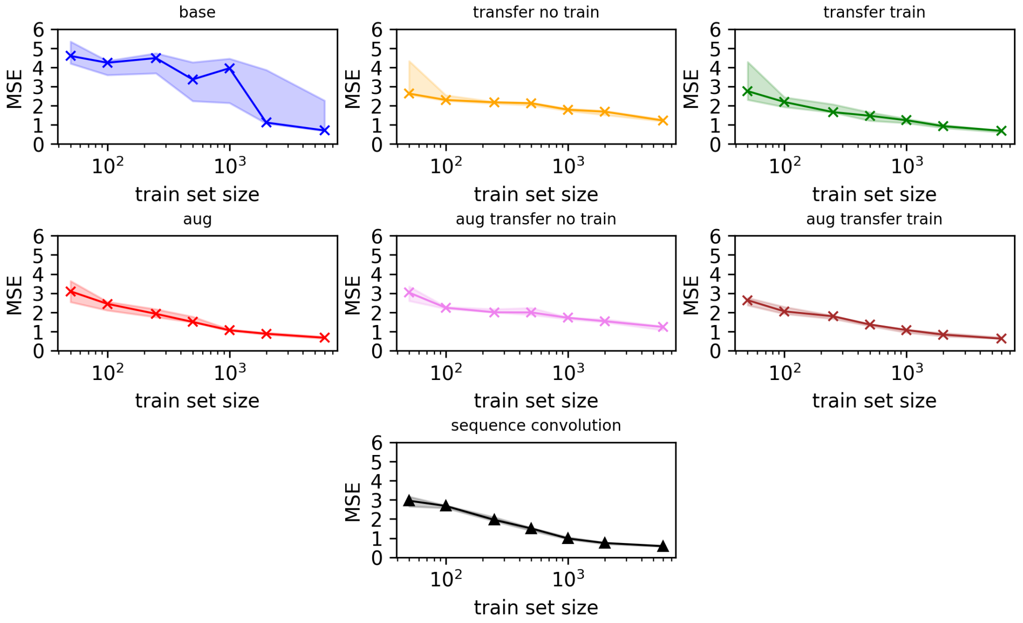


Figure S20: Spread of MSE values of the different training runs for DenseNet trained on Pab1

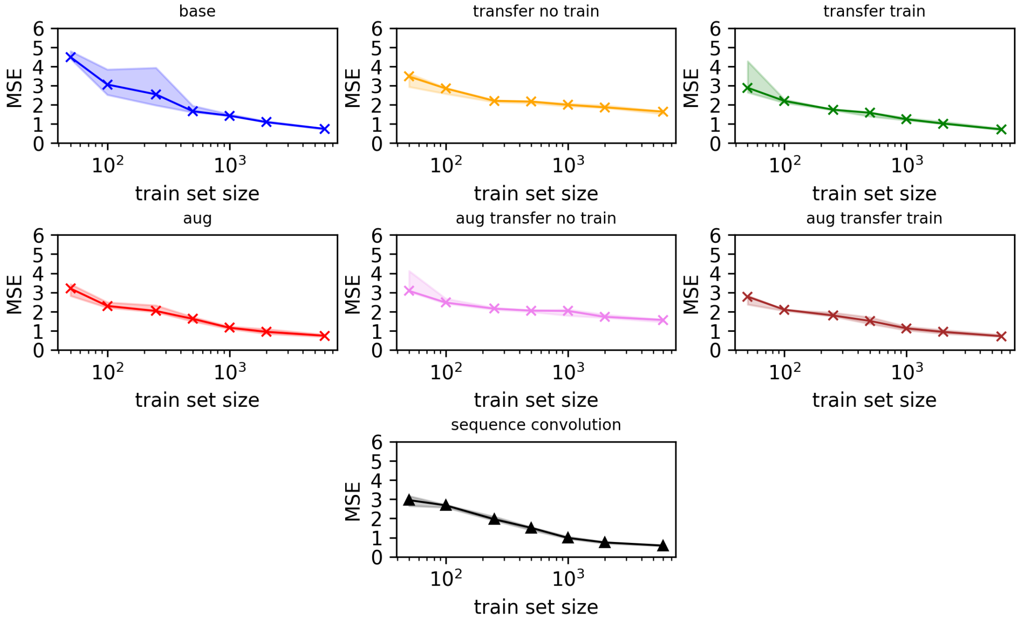


Figure S21: Spread of MSE values of the different training runs for SepConvMixer trained on Pab1

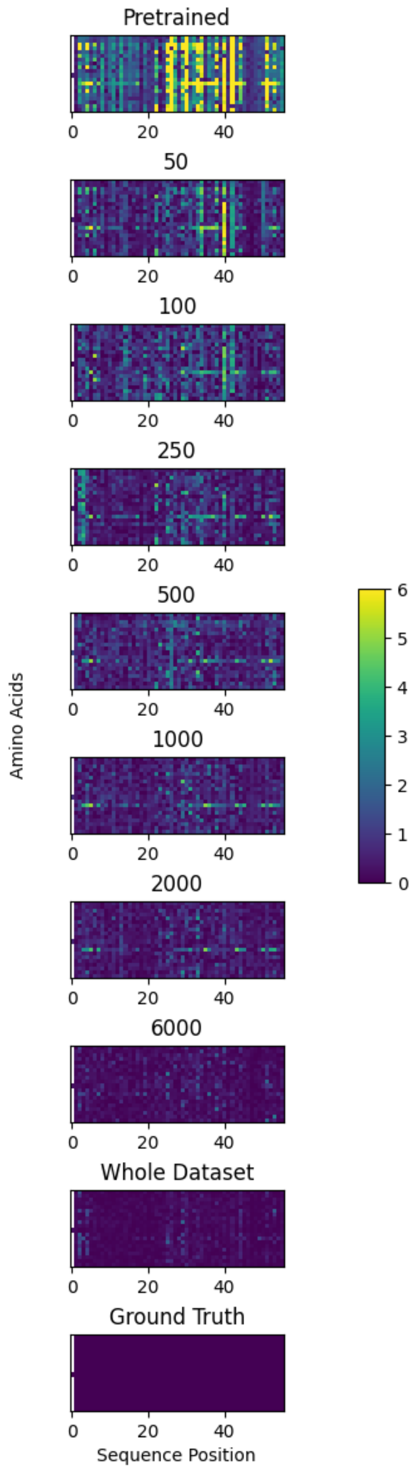


Figure S22: Heatmaps showing the absolute difference of the predictions to the ground truth

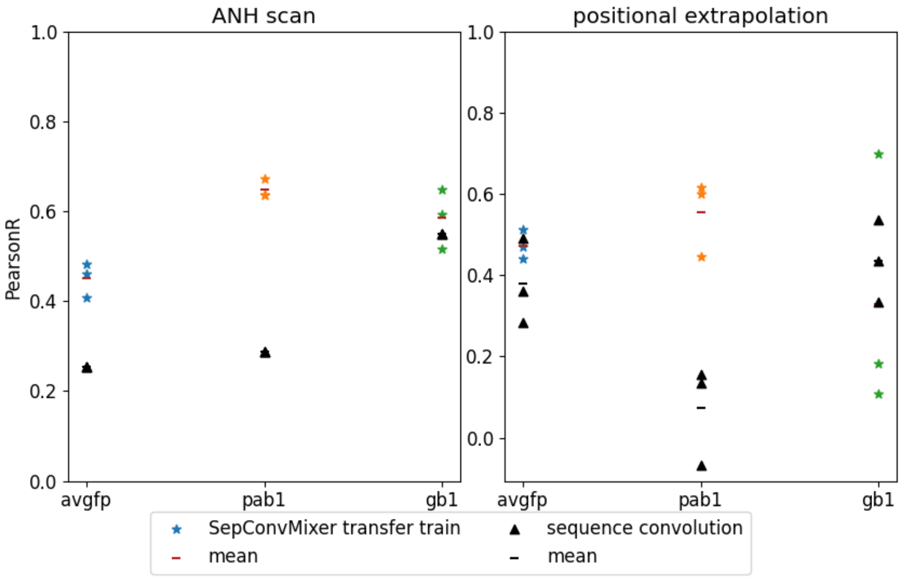


Figure S23: PearsonR for predictions on an ANH-Scan as well as on positions the networks (pre-trained SepConvMixer and sequence convolution) have not seen before
.

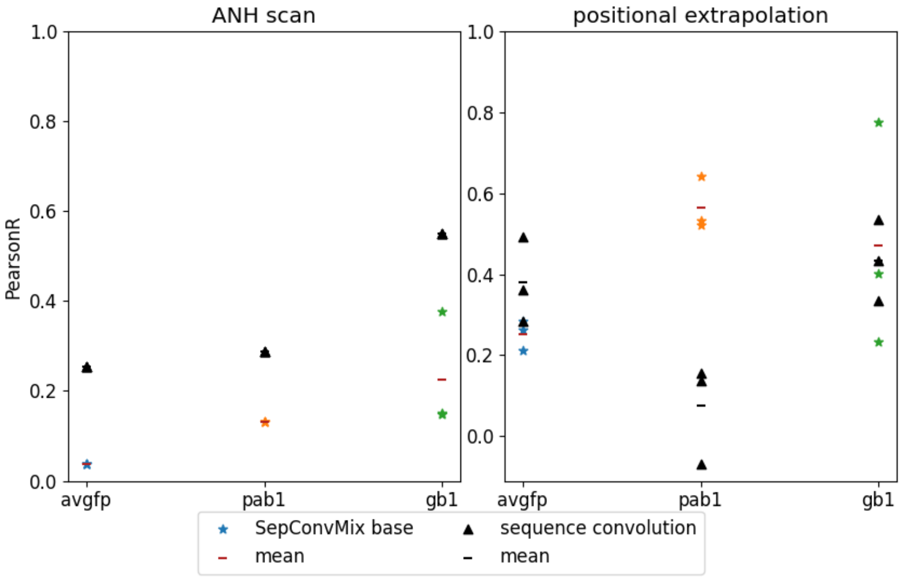


Figure S24: PearsonR for predictions on an ANH-Scan as well as on positions the networks (SepConvMixer and sequence convolution) have not seen before
.

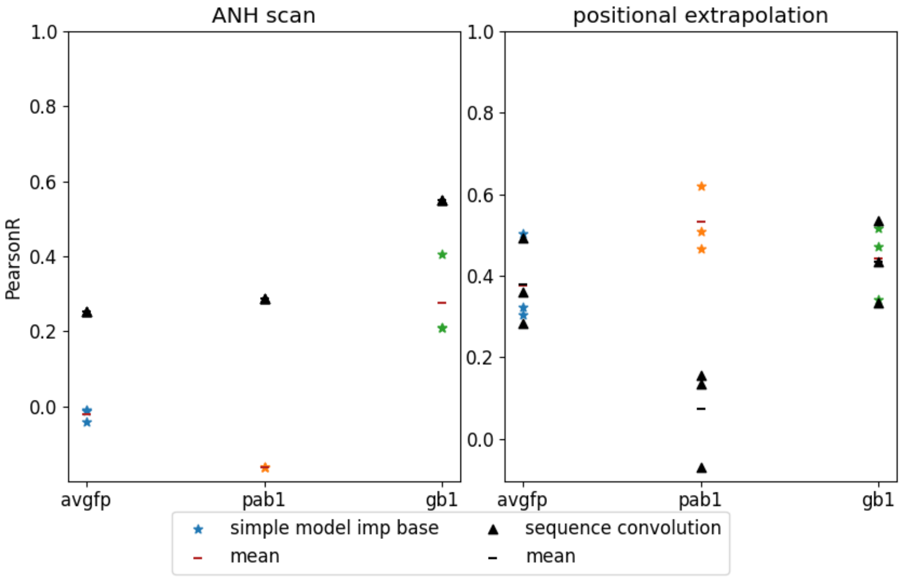


Figure S25: PearsonR for predictions on an ANH-Scan as well as on positions the networks (simple CNN and sequence convolution) have not seen before
.
